# Inhibition of HIV infection by structural proteins of the inner nuclear membrane is associated with reduced chromatin dynamics

**DOI:** 10.1101/2020.12.03.410522

**Authors:** Anvita Bhargava, Mathieu Maurin, Patricia M. Davidson, Mabel Jouve, Xavier Lahaye, Nicolas Manel

**Affiliations:** Institut Curie, PSL Research University, INSERM U932, Paris, France; Laboratoire Physico-Chimie Curie, Institut Curie, CNRS UMR168, Sorbonne Université, PSL Research University, Paris, France; Institute Curie, UMR3215, Paris, France

## Abstract

The Human Immunodeficiency Virus (HIV) enters the nucleus to establish infection. HIV interacts with nuclear pore components to cross the nuclear envelope. In contrast, the role of other proteins of the nuclear envelope in HIV infection is not yet understood. The inner nuclear transmembrane proteins SUN1 and SUN2 connect lamins in the interior of the nucleus to the cytoskeleton in the cytoplasm. Increased levels of SUN1 or SUN2 potently restrict HIV infection through an unresolved mechanism. Here, we find that SUN1 and SUN2 exhibit a differential and viral strain-specific antiviral activity HIV-1 and HIV-2. In macrophages and HeLa cells, HIV-1 and HIV-2 are respectively preferentially inhibited by SUN1 and SUN2. This specificity maps to the nucleoplasmic domain of SUN proteins, which associates with Lamin A/C and participates to the DNA damage response. We find that etoposide, a DNA-damaging drug, stimulates infection. Inhibition of the DNA damage signaling kinase ATR, which induces a DNA damage response, also enhances HIV-1 infection. The proviral effect of ATR inhibition on infection requires the HIV-1 Vpr gene. Depletion of endogenous Lamin A/C, which sensitizes cells to DNA damage, also enhances HIV-1 infection in HeLa cells. SUN1 overexpression neutralizes these proviral effects, while the antiviral effect of SUN2 is rescued by etoposide treatment. Finally, we show that inhibition of HIV-1 infection by overexpressed SUN proteins and endogenous Lamin A/C is associated with reduced internal movements of chromatin and reduced rotations of the nucleus. Altogether, these results highlight distinct antiviral activities of SUN1 and SUN2 and reveal an emerging role of nuclear movements and the DNA damage response in the control of HIV infection by structural components of the nuclear envelope.

## Introduction

Successful infection of cells by HIV requires an active transport of the virus through the physical barrier of the nuclear envelope. Nuclear entry of HIV is coordinated with the completion of reverse transcription and selection of integration sites (Dharan et al., 2020; Schaller et al., 2011). The capsid protein of HIV engages multiple interactions with nuclear pore complex (NPC) components and associated proteins such as Cyclophilin A to achieve this coordination (Yamashita and Engelman, 2017).

In the nuclear envelope, in addition to NPC proteins, SUN proteins located at the inner nuclear membrane impact HIV infection (Bhargava et al., 2018). SUN1 and SUN2 are integral proteins of the inner nuclear envelope of somatic cells. They play essential roles in the maintenance of genomic stability and the resolution of DNA damage (Lawrence et al., 2016; Lei et al., 2012). SUN proteins possess a lamin-binding domain at their N-terminus located in the nucleoplasm. Lamins are intermediated filament proteins that assemble the nuclear lamina, a dense meshwork contributing to mechanical protection, organization of chromatin domains and recruitment of DNA repair factors (Burke and Stewart, 2013; Gonzalo, 2014). At their C-terminus, SUN proteins interact with the KASH domains of nesprins in the perinuclear space. Nesprins are large integral proteins of the outer nuclear membrane (Burke and Stewart, 2013). Nesprins have multiple interactions with cytoskeletal proteins, enabling a dynamic anchoring of the nucleus within the cells.

SUN2 was first identified as an antiviral factor against HIV-1 in the context of a cDNA screen (Schoggins et al., 2011). Subsequent studies confirmed and extended the antiviral viral effect of SUN1 and SUN2 overexpression on HIV-1 and HIV-2 infection (Donahue et al., 2016; Lahaye et al., 2016; Luo et al., 2018; Schaller et al., 2017). SUN1 and SUN2 overexpression limits the level of HIV-1 nuclear import (Donahue et al., 2016; Luo et al., 2018; Schaller et al., 2017), leading to reduced viral integration. Furthermore, nanotubes of HIV-1 capsid and nucleocapsid proteins produced *in vitro*, pull down SUN1 and SUN2 proteins from cell lysates, suggesting that SUN proteins and the viral capsid protein may interact directly or indirectly during infection (Schaller et al., 2017).

The role of endogenous SUN2 in HIV-1 infection has been examined but a consensus has not been reached (Donahue et al., 2017; Lahaye et al., 2016; Schaller et al., 2017; Sun et al., 2018). Three studies concurred with a requirement for SUN2 in HIV-1 infection in primary CD4+ T cells, in monocyte-derived dendritic cells and in THP-1 cells, although the strength of this requirement varies between cell type (Donahue et al., 2017; Lahaye et al., 2016; Schaller et al., 2017). A fourth study obtained contradicting results and proposed that endogenous SUN2 instead limits HIV infection at the level of viral promoter expression (Sun et al., 2018). We initially proposed that HIV infection requires an optimal level of SUN2 protein, and that both depletion and overexpression impair infection, not necessarily through the same mechanism (Lahaye et al., 2016). This notion fits well with the structural role of the LINC complex in nuclear architecture. Of note, endogenous SUN2 level varies with the extent of T cell activation (Sun et al., 2018). It is thus conceivable that variable experimental conditions between studies, particularly using sensitive primary immune cells, could account for the variable effects of endogenous SUN2 on HIV infection. SUN2 is also implicated in the effects of Cyclophilin A (CypA) on HIV-1 infection. In HeLa cells, SUN2 overexpression abrogates the sensitivity of HIV-1 capsid mutant N74D to Cyclophilin A inhibition (Lahaye et al., 2016). In primary CD4+ T cells and murine bone-marrow derived dendritic cells, endogenous SUN2 is required for the Cyclophilin A-dependent steps of HIV infection (Lahaye et al., 2016). Another study however, did not observe this effect in primary CD4+ T cells (Donahue et al., 2017). These differences may reflect the use of different read-outs for quantifying the impact of cyclophilin A inhibition on infection.

Our understanding of the antiviral effect of SUN1 is less advanced. In HEK293A cells, the antiviral effect of SUN1 overexpression requires the interaction of Cyclophilin A with HIV-1 capsid protein (Luo et al., 2018). In THP-1 cells, endogenous SUN1 is not required for HIV-1 infection (Schaller et al., 2017).

The strong antiviral effect of SUN protein overexpression on HIV infection exploits one or several points of weakness in the viral replication cycle. The cellular mechanisms by which elevated levels of SUN expression block HIV infection are not known. Intriguingly, SUN2 overexpression is associated with alteration of nuclear envelope shape, suggesting that SUN might interfere with HIV infection through a perturbation of the integrity of the nucleus (Donahue et al., 2016; Lahaye et al., 2016). However, it has not been possible so far to explain how SUN proteins are perturbing cellular and nuclear physiology to impact HIV.

## Results

### SUN1 and SUN2 proteins demonstrate HIV strain-specific antiviral effects

To gain insights in SUN1- and SUN2-mediated antiviral effects on the early steps of HIV infection, we first performed a comparative assessment of the antiviral effect of SUN1 and SUN2 on HIV infection in primary cells. To this end, we overexpressed SUN1 and SUN2 in primary monocyte-derived macrophages (MDMs) using lentiviral vectors (**Figure 1A**). In order to focus on the early phase of infection, cells were infected using single-round HIV-1 and HIV-2 encoding GFP in the place of the Nef gene. SUN1 and SUN2 induced an antiviral effect on HIV-1 and HIV-2 (**Figure 1B**). Unexpectedly, SUN1 and SUN2 did not show an identical antiviral effect on the two strains. The calculation of the ratio of inhibition by SUN1 over SUN2 revealed that HIV-1 was preferentially inhibited by SUN1, while HIV-2 was preferentially inhibited by SUN2 (**Figure 1B**). In MDMs, HIV-1 infection is sensitive to inhibition by Cyclosporin A (CsA). CsA treatment did not further inhibit infection in cells overexpressing SUN1 or SUN2 (**Figure 1B**). We analyzed the progression of HIV-1 and HIV-2 infection in the context of SUN protein expression using RT-qPCR on viral DNA species. For HIV-1, both SUN1 and SUN2 overexpression reduced the level of integrated viral DNA (**Figure 1C**). However, SUN1 reduced the total amount of viral DNA, while SUN2 reduced the level of 2-LTR circles, that are a hallmark of viral entry into the nucleus. CsA reduced the total amount of viral DNA in control cells and there was no additional reduction following SUN protein expression. For HIV-2, SUN1 and SUN2 significantly reduced 2-LTR circles only. These experiments indicate that SUN1 and SUN2 have strain-specific antiviral effects and that they modify different steps of viral infection.

**Figure 1.**
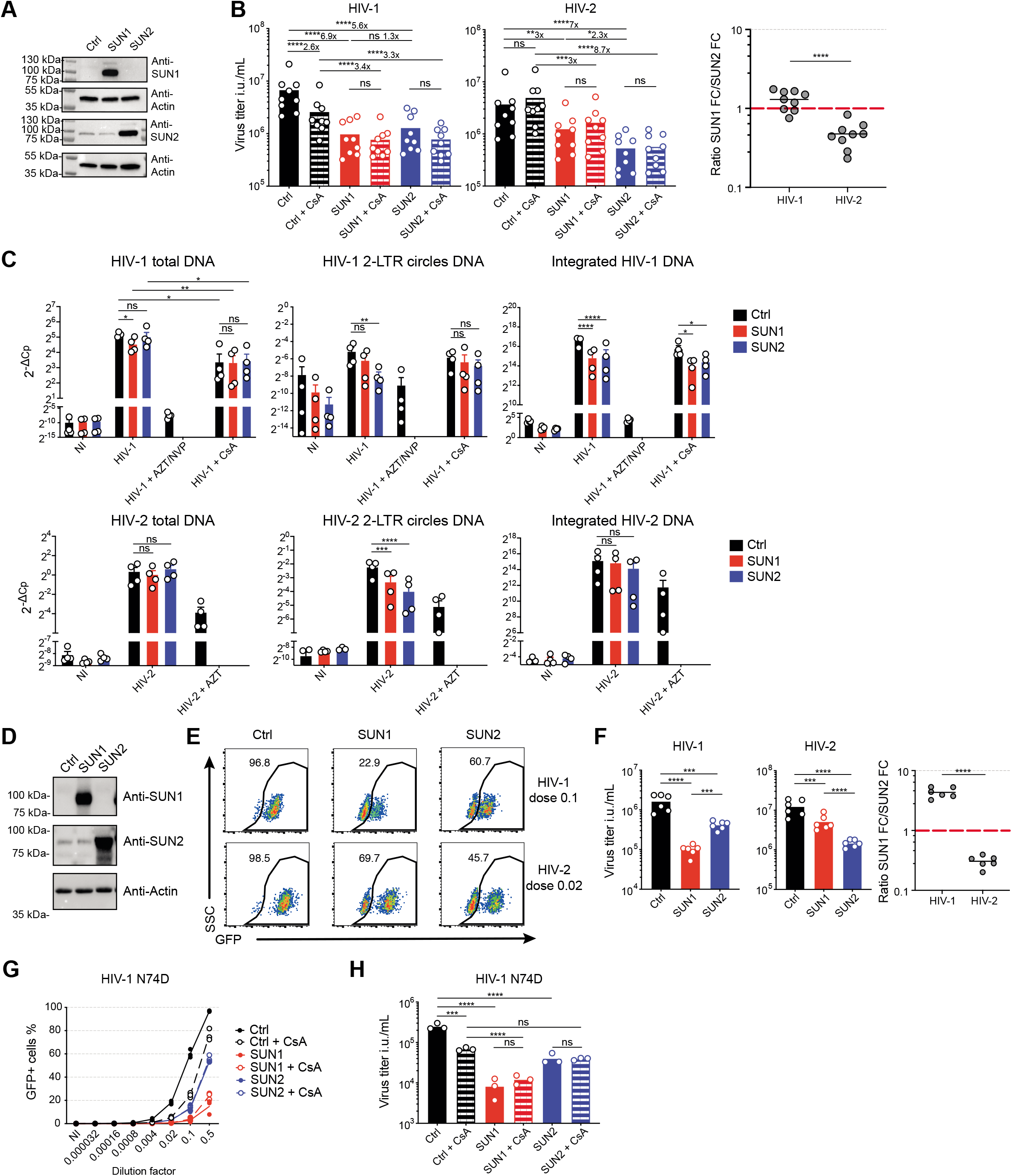
Distinct antiviral activities of SUN1 and SUN2 against HIV-1 and HIV-2. **(A)** Detection of SUN1, SUN2 and actin in MDMs transduced with mTagBFP-2A control, SUN1 or SUN2 lentivectors (representative of n = 3). **(B)** Left, viral titers as infectious units (i.u.) per mL based on percentages of GFP^+^ MDMs 48 hours after infection with serial dilutions of HIV-1 or HIV-2 encoding GFP in Nef and pseudotyped with VSV-G, with or without 2 μM CsA (n = 9 donors, paired RM ANOVA one-way on Log-transformed titers with Sidak post-test, line at mean). Right, ratios of titer fold changes (FC) control over SUN1 (SUN1 FC) or control over SUN2 (SUN2 FC) (paired t-test, line at mean). **(C)** Detection of HIV-1 total DNA, 2-LTR circles DNA and integrated DNA by RT-qPCR at 24 hours after infection with HIV-1 or HIV-2 (dilution factor: 0.17) of MDMs transduced with mTagBFP-2A control, SUN1 or SUN2 lentivectors. Reverse transcriptase inhibitors Azidothymidine (AZT; 24 μM) and Nevirapine (NVP; 10 μM) were added during infection only on control cells (n=4 donors, paired RM ANOVA one-way with Sidak post-test, line at mean ± SEM). **(D)** Detection of SUN1, SUN2 and actin in HeLa cells transduced with mTagBFP-2A control, SUN1 or SUN2 lentivectors. **(E)** GFP expression in BFP-positive HeLa cells transduced with mTagBFP-2A control, SUN1 or SUN2 lentivectors, 48 hours after infection with indicated dilutions of HIV-1 and HIV-2 (representative data from one experiment at the indicated dose of virus). **(F)** Left, Viral titers based on percentages of GFP^+^ cells after infection with serial dilutions of virus as in **(E)** (n = 6, paired RM ANOVA one-way on Log-transformed titers, with Dunnet’s post-test, line at mean). Right, ratios calculated as in **(B)**. **(G)** Percentage of GFP^+^ in BFP^+^ HeLa cells transduced with mTagBFP-2A control, SUN1 or SUN2 lentivectors, 48 hours after infection with serial dilutions of HIV-1 or HIV-1 CA N74D, with or without treatment with 2 μM of CsA (n = 3 independent experiments). **(H)** Viral titers as in (G) (n = 3, paired RM ANOVA one-way on Log-transformed titers with Sidak’s post-test, line at mean). Ctrl = control, *p < 0.05, ***p < 0.001, ****p < 0.0001; ns, not statistically significant.

We similarly overexpressed SUN1 and SUN2 in HeLa cells (**Figure 1D**). SUN1 overexpression had a greater inhibitory effect on HIV-1 infection than SUN2 overexpression, whereas in HIV-2 infection, SUN2 overexpression had a greater effect than SUN1, recapitulating the results obtained in MDMs (**Figure 1E, 1F**). In HeLa cells, wild-type HIV-1 is not sensitive to CsA, but HIV-1 CA N74D is, similar to HIV-1 WT in MDM (De Iaco and Luban, 2014). We thus used this mutant to address the relationship between the antiviral effect of SUN and CsA sensitivity. Both SUN1 and SUN2 abolished CsA sensitivity of HIV-1 CA N74D in HeLa cells (**Figure 1G, 1H**). We next measured the levels of HIV-1 DNA species. SUN1 and SUN2 reduced the levels of integrated HIV-1 DNA and this effect was more pronounced for SUN1 (**Figure S1A**). Similar to MDM, SUN1 significantly inhibited the level of total viral DNA while the levels of 2-LTR circles were not significantly reduced. In contrast, SUN2 did not impact the level of total viral DNA but reduced the level of 2-LTR circles. We thus focused on HeLa cells for additional experiments aiming at characterizing the strain-specific inhibition of SUN1 and SUN2.

### Strain-specific antiviral activity maps to the nucleoplasmic domain of SUN proteins

Cell-cell communication factors of innate immunity, such as interferons and cGAMP, can contribute to antiviral effects on top of cell-intrinsic restriction factors. Using a co-culture of SUN1/2-expressing cells and control cells expressing a fluorescent marker (TagRFP657), we found that the strain-specific effect of SUN1 and SUN2 on HIV-1 and HIV-2 infection is entirely cell-intrinsic in HeLa cells (**Figure 2A**). To determine if SUN1 and SUN2 induced an antiviral state at the cell-intrinsic level through expression of other antiviral genes, we performed a transcriptomic analysis of SUN1 and SUN2 overexpressing cells. Strikingly, we could not detect any differentially expressed gene in this dataset, aside from SUN1 and SUN2 themselves (**Figure S1B**). Next, we generated chimeras between SUN1 and SUN2 to map the strain-specific antiviral effect (**Figure 2B, 2C**). We found that the N-terminal nucleoplasmic domains of SUN1 and SUN2 confer strain-specificity (**Figure 2D, 2E**). These results establish that SUN1 and SUN2 exert a cell-intrinsic HIV-strain specific antiviral effect on HIV infection, that maps to the nucleoplasmic domain of SUN proteins.

**Figure 2.**
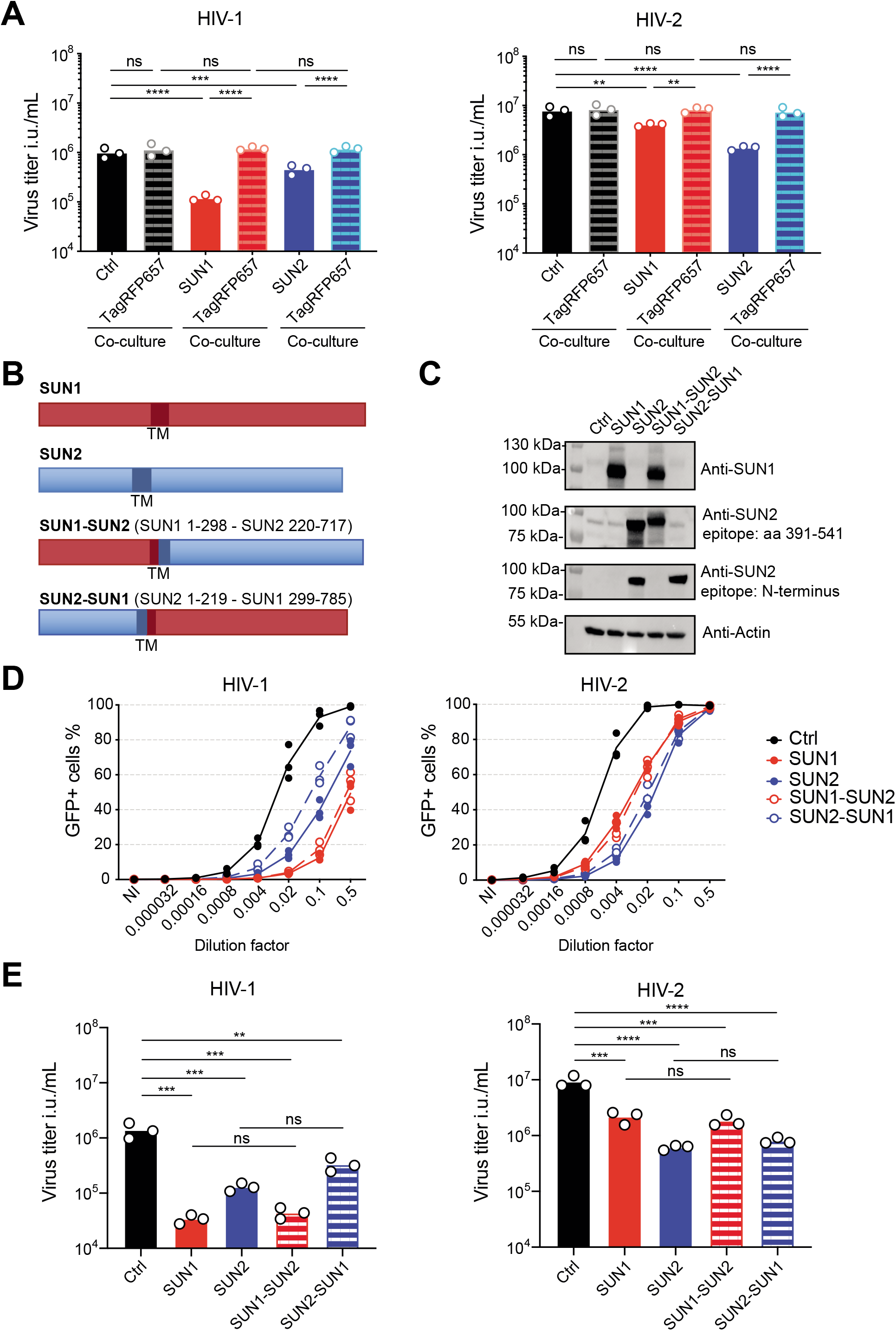
Mapping of the strain-specific antiviral activity of SUN proteins. **(A)** mTagBFP-2A Ctrl, mTagBFP-2A-SUN1 and mTagBFP-2A-SUN2 expressing HeLa cells were co-cultured at a 1:1 ratio with HeLa cells expressing TagRFP657-2A and infected with serial dilutions of HIV-1 and HIV-2. Titers were calculated based on percentage of GFP^+^ cells 48 hours post-infection dilutions within the indicated populations (n=3 independent experiments, paired RM ANOVA one-way on Log-transformed titers with Sidak’s post-test, line at mean). **(B)** Schematic representation of chimeric proteins between full-length SUN1 (red) and SUN2 (blue). Amino-acid residues retained in hybrid proteins are indicated within brackets. **(C)** Detection of SUN1, SUN2 and actin in HeLa cells transduced with the indicated mTagBFP-2A lentivectors. Two antibodies targeting SUN2 that recognize different epitopes within the protein were used. **(D)** Percentage of GFP^+^ in BFP^+^ HeLa cells transduced with the indicated mTagBFP-2A lentivectors, 48 hours after infection with serial dilutions of HIV-1 or HIV-2 (n=3 independent experiments). **(E)** Viral titers based on percentages of GFP^+^ cells shown in **(D)** (n=3, paired RM ANOVA one-way with Sidak’s post-test, line at mean). Ctrl = control, *p < 0.05, **p < 0.01, ***p < 0.001, ****p < 0.0001; ns, not statistically significant.

### Antiviral effect of SUN proteins at the nuclear envelope

We next sought to study how SUN1 and SUN2 impact cells to inhibit HIV infection. Electron microscopy analysis revealed that both SUN1 and SUN2 overexpression induced deep invaginations of the nuclear envelope, that appeared more pronounced with SUN2 (**Figure 3A**). This raised the possibility that alteration of the shape of the nucleus could be responsible for the antiviral effect. To test this, we asked if the antiviral effect of SUN occurs at the nuclear envelope or whether it is the result of cytosolic accumulation of SUN proteins. SUN proteins form the LINC complex with nesprins at the nuclear envelope by interaction with their KASH domain within the perinuclear space. Expression of the isolated KASH domain (spectrin repeat-KASH, SR-KASH) functions as a dominant negative by disrupting the SUN-Nesprin interaction and displacing Nesprins from the nuclear envelope (Starr et al., 2003). We co-expressed SR-KASH with SUN proteins (**Figure 3B**). We found that co-expression of the SR-KASH construct partially rescued the antiviral effect on SUN1 and SUN2 on HIV-1 infection (**Figure 3C)**. These results support the notion that SUN proteins have an antiviral activity at the nuclear envelope.

**Figure 3.**
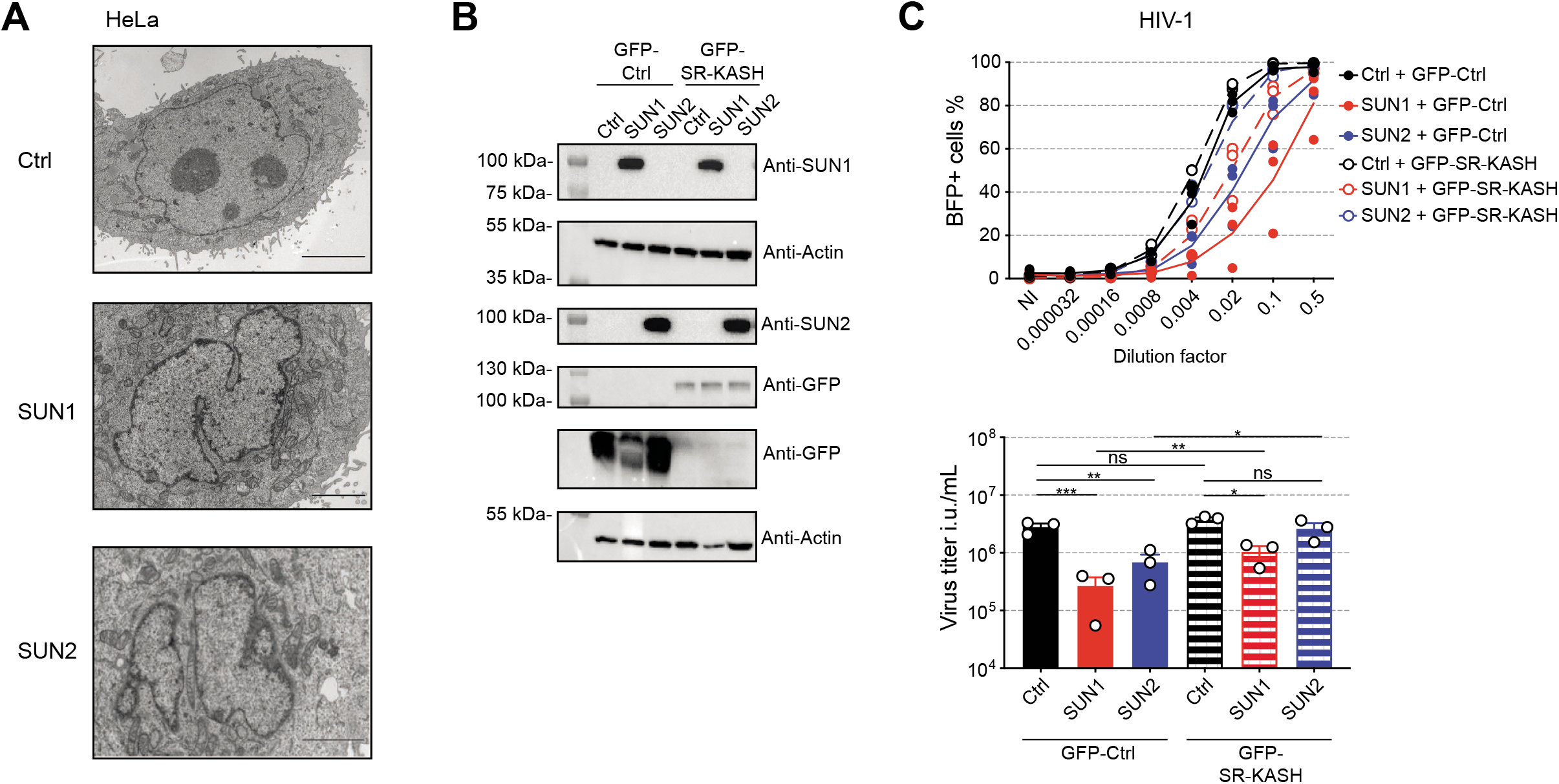
SUN proteins inhibit HIV infection at the nuclear envelope. **(A)** Representative electron micrograph showing nuclei in control, SUN1 and SUN2 overexpressing HeLa cells (scale: 10, 2 and 5 μm respectively). **(B)** Detection of SUN1, SUN2, GFP and actin in HeLa cells transduced with TagRFP657-expressing control, SUN1 or SUN2 lentivectors, combined with control GFP or SR-KASH DN fused to GFP expressing lentivectors. The same lysates were loaded onto two separate membranes, the housekeeping control is shown for both. **(C)** Top, percentage of BFP^+^ within tagRFP657^+^ HeLa cells 48 hours after infection with serial dilutions of HIV-1 encoding BFP in the place of Nef, pseudotyped with VSV-G. Bottom, viral titers based on percentages of BFP^+^ cells (n=3, paired RM ANOVA one-way on Log-transformed titers, with Sidak’s post-test, line at mean ± SEM). Ctrl = control, *p < 0.05, **p < 0.01, ***p < 0.001; ns, not statistically significant.

### Interplay between SUN proteins, HIV infection and the DNA damage response

As a next step, we attempted to characterize the cellular processes affected by elevated levels of SUN proteins. SUN1 and SUN2 are required to limit the accumulation of DNA damage in cells (Lei et al., 2012). Since HIV infection is a DNA-damaging event, we considered the possible interplay between SUN proteins, HIV infection and the DNA damage response. We examined the level of γH2AX, an early marker of the DNA damage response. At baseline, we did not detect any modification of γH2AX levels upon SUN1 or SUN2 expression (**Figure 4A, 4B**). To induce DNA damage, we selected etoposide, a topo-isomerase II inhibitor. Interestingly, SUN1 overexpression but not SUN2, significantly limited the levels of induced γH2AX after etoposide treatment (**Figure 4A, 4B**). To explore the potential link between DNA damage and infection, we infected HeLa cells in the presence of etoposide for the first 4 hours of the experiment. However, etoposide gradually induces apoptosis of treated cells (Rello-Varona et al., 2006), which hampered our ability to detect viable cells to measure infection after 48 hours. To circumvent this, we cultured cells in the presence of the caspase inhibitor Q-VD-Oph and lower doses of etoposide. Q-VD-Oph did not prevent γH2AX induction by etoposide treatment (**Figure 4C**). Etoposide treatment increased HIV-1 infection by 2-fold on average (**Figure 4D**). HIV-2 infection was also increased but significantly at only one dose of etoposide tested, with a smaller fold change. Next, we combined SUN expression with etoposide to determine the epistatic relationship between DNA damage induction and SUN expression on the level of HIV infection. Here, we used a higher MOI to observe the antiviral effect of SUN proteins. The increase induced by etoposide treatment on control cells was consistently observed across experiments. Interestingly, SUN1 abrogated the proviral effect of etoposide treatment and in sharp contrast, etoposide treatment rescued cells from the antiviral effect of SUN2 (**Figure 4E, 4F**). Thus, SUN1 overexpression, which restricts HIV-1 infection more than SUN2, operates downstream of DNA damage induction, while SUN2 overexpression impacts infection upstream of DNA damage.

**Figure 4.**
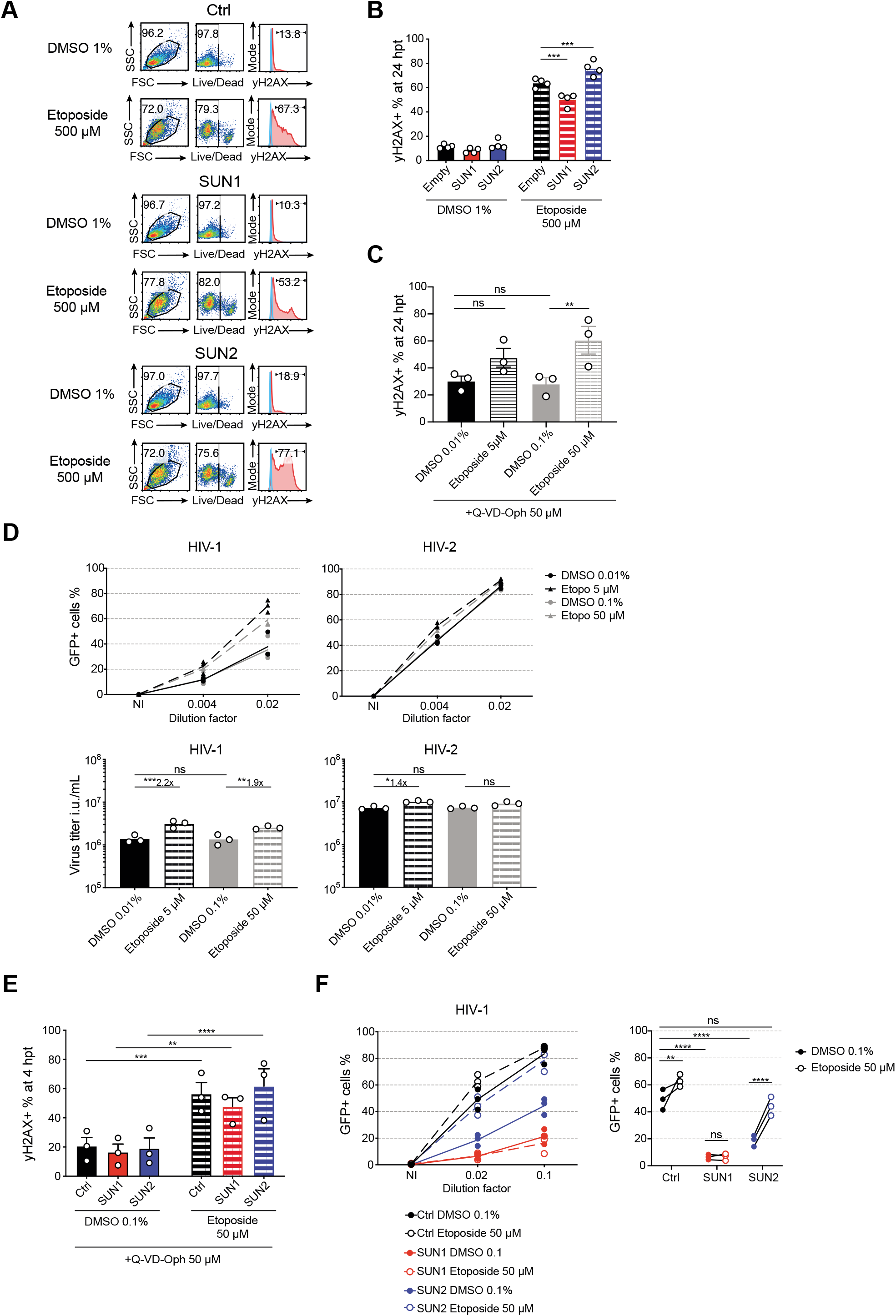
Interplay between HIV-1 infection, SUN protein and the DNA damage response. **(A)** Viability and gH2AX intracellular staining in HeLa cells transduced with mTagBFP-2A control, SUN1 or SUN2 lentivector, 24 hours after treatment with 500 μM of etoposide or 1% DMSO as control (representative experiment from n = 3). **(B)** Quantification of gH2AX^+^ HeLa cells treated as in **(A)** (n=4, paired RM ANOVA one-way with Sidak’s post-test, line at mean). **(C)** Quantification of gH2AX^+^ HeLa cells 24 hours after a 4-hour treatment with 5 μM, 50 μM etoposide or corresponding DMSO control. 50 μM of Q-VD-Oph were present throughout the experiment (n=3, paired RM ANOVA one-way with Sidak’s post-test, line at mean ± SEM). **(D)** Top, percentage of GFP^+^ HeLa cells 48 hours after infection with two dilutions of HIV-1 or HIV-2, treated as in **(C)**. Cells were treated and infected simultaneously, the drugs and the virus were washed out at 4 hours post-treatment/infection. 50 μM of Q-VD-Oph were maintained throughout the experiment. Bottom, viral titers based on percentages of GFP^+^ cells (n=3, paired RM ANOVA one-way with Sidak’s post-test, line at mean). **(E)** Quantification of γH2AX^+^ HeLa cells, transduced with mTagBFP-2A control, SUN1 or SUN2 lentivectors, 4 hours after treatment with 50 μM of etoposide or corresponding DMSO control (n=3, paired RM ANOVA one-way with Sidak’s post-test, line at mean ± SEM). **(F)** Left, percentage of GFP^+^ cells in BFP^+^ HeLa cells expressing control, SUN1, SUN2 lentivectors and treated as in **(E)**, 48 hours after infection with purified HIV-1 env-nef-, expressing GFP in the place of Nef and pseudotyped with VSV-G. Right, GFP+ percentages at viral dilution 0.02 (n=3, experimental pairs are indicated; RM ANOVA two-way test, uncorrected Fischer’ LSD). 50 μM of Q-VD-Oph were maintained throughout the experiment. Ctrl = control, hpt = hours post treatment, *p < 0.05, **p < 0.01, ***p < 0.001, ****p < 0.0001; ns, not statistically significant.

### DNA damage induction by ATR inhibition and role of HIV-1 Vpr

Next, we looked for a different approach to induce DNA damage that would be functionally linked to the nuclear envelope. ATR is a DNA damage sensor and functions as a checkpoint at the nuclear envelope in response to mechanical stress (Kumar et al., 2014). ATR inhibition heightens DNA damage in cells (Foote et al., 2018). Furthermore, in HIV-1, expression of the accessory protein Vpr causes DNA damage and activates ATR (Roshal et al., 2003), although the relevance of this effect in the context of virion-packaged Vpr is unknown. Considering the significance of ATR at the nuclear envelope and its relationship with Vpr, we asked if DNA damage induction by ATR inhibition, SUN and Vpr are functionally related. We inhibited ATR using AZD6738, a next-generation inhibitor with improved specificity (Foote et al., 2018). As expected, ATR inhibition increased the levels of γH2AX in HeLa cells (**Figure 5A**). We next infected HeLa cells with p24-normalized and sucrose-cushion purified stocks of the HIV-1 single-round virus and its HIV-1 vpr-deficient (vpr-) counterpart. ATR inhibition increased vpr-positive HIV-1 infection in HeLa cells by 2-fold, similar to etoposide treatment (**Figure 5B, left panel**). Concomitant SUN protein overexpression inhibited HIV-1 infection and abrogated the proviral effect of ATR inhibition, again similar to the etoposide treatment. Unexpectedly, the titer of the p24-normalized HIV-1 vpr- was slightly higher than the HIV-1 wild-type counterpart in HeLa cells, and HIV-1 vpr- was insensitive to ATRi (**Figure 5B, right panel**). However, HIV-1 vpr- remained fully sensitive to the antiviral effect of SUN protein expression. In sum, this data supports the idea that increased DNA damage favors HIV-1 infection of HeLa cells and that SUN1 operates downstream of this to block infection. Intriguingly, this data also revealed that sensitivity to ATR inhibition is a HIV-1 vpr phenotype in single-round infection of HeLa cells.

**Figure 5.**
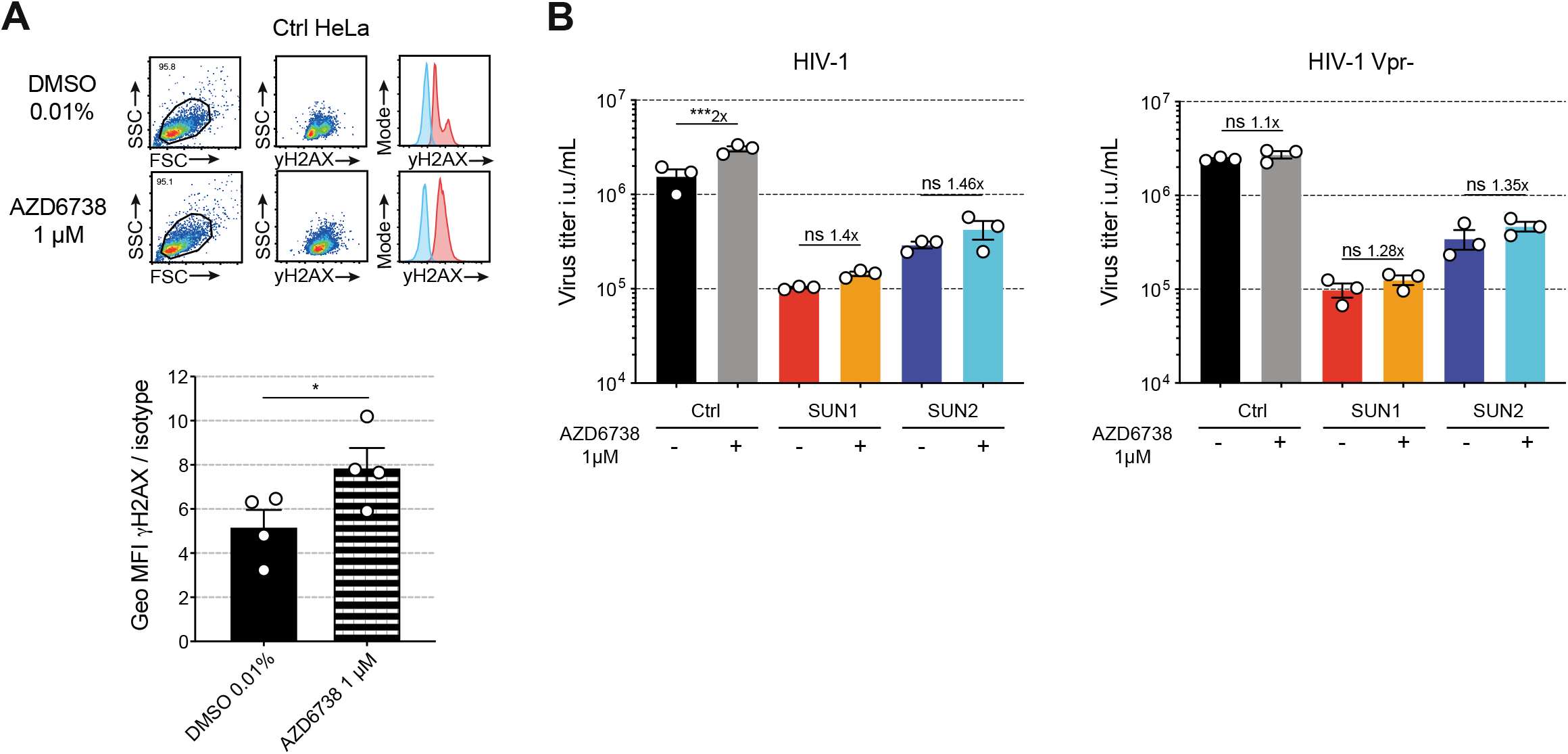
DNA damage induction by ATR inhibition stimulates HIV-1 infection in a Vpr- dependent manner. **(A)** Top, γH2AX intracellular staining in mTagBFP-2A control HeLa cell line, 24 hours after treatment with either 1 μM of AZD6738 or 0.01% DMSO (representative experiment). Bottom, quantification of specific γH2AX staining as geometric mean fluorescence intensity (GeoMFI) of the antibody signal divided by the GeoMFI of the isotype control (n=4, paired t-test, *p < 0.05, line at mean ± SEM). **(B)** Viral titers based on percentages of GFP^+^ cells, within BFP^+^ HeLa cells expressing Ctrl, SUN1 or SUN2 lentivector, with or without treatment with 1 μM of AZ6738, 48 hours after infection with serial dilutions of purified, p24-normalized, HIV-1 env-nef- or HIV-1 env-nef-vpr- encoding GFP in the place of Nef, pseudotyped with VSV-G (n=3, paired RM ANOVA one-way with Sidak’s post-test, line at mean ± SEM). Ctrl = control, hpt = hours post treatment, *p < 0.05, ***p < 0.001, ns, not statistically significant.

### Endogenous Lamin A/C limits HIV-1 infection in HeLa cells

We sought to further explore the relationship between infection, DNA damage and structure of the nuclear envelope. We searched for an orthogonal approach to perturb the nuclear envelope structure and the DNA damage response. Lamin A/C expression is required to maintain a regular nuclear shape (Lammerding et al., 2004) and to protect from DNA damage (Singh et al., 2013). We used short-hairpin RNA to reduce expression of Lamin A/C (**Figure 6A**). Similar to SUN protein overexpression, knock-down of Lamin A/C compromised the regularity of the nuclear envelope shape (**Figure 6B**). To quantify this effect, we measured the shape descriptor ‘solidity’ of the nucleus: solidity values close to 1 indicate smoothly convex nuclei while lower values correspond to deformed, lobulated nuclei, presenting concave invaginations. Overexpression of SUN proteins and silencing of lamin A/C increased nuclear envelope deformation, although this effect was less pronounced in cells overexpressing SUN1 (**Figure 6C**). The endogenous levels of Lamin A/C and SUN proteins were not reciprocally affected by SUN overexpression or Lamin A/C silencing. (**Figure 6A**). After infection, Lamin A/C knock-down unexpectedly increased HIV-1 infection levels by 1.6-fold, while HIV-2 infection was not affected (**Figure 6D**). Lamin A/C thus limits HIV-1 infection in HeLa cells. However, the increase in nuclear envelope shape irregularities does not explain how lamin A/C depletion and SUN protein overexpression affect HIV infection. Given the opposing effects of SUN overexpression and Lamin A/C knock-down on HIV-1 infection, we examined if the antiviral effect of SUN proteins requires endogenous lamins. We knocked-down Lamin A/C, Lamin B1 and Lamin B2 and co-expressed SUN proteins (**Figure 6E**). Viable Lamin B1-depleted HeLa cells could not be maintained in culture. Lamin A/C depletion enhanced HIV-1 infection as above, and Lamin B2 depletion had no effect (**Figure 6F**). SUN proteins maintained their antiviral effect irrespective of the level of Lamin A/C and Lamin B2. This shows that the effect of elevated levels of SUN proteins is dominant on the effect of Lamin A/C depletion on HIV-1 infection.

**Figure 6.**
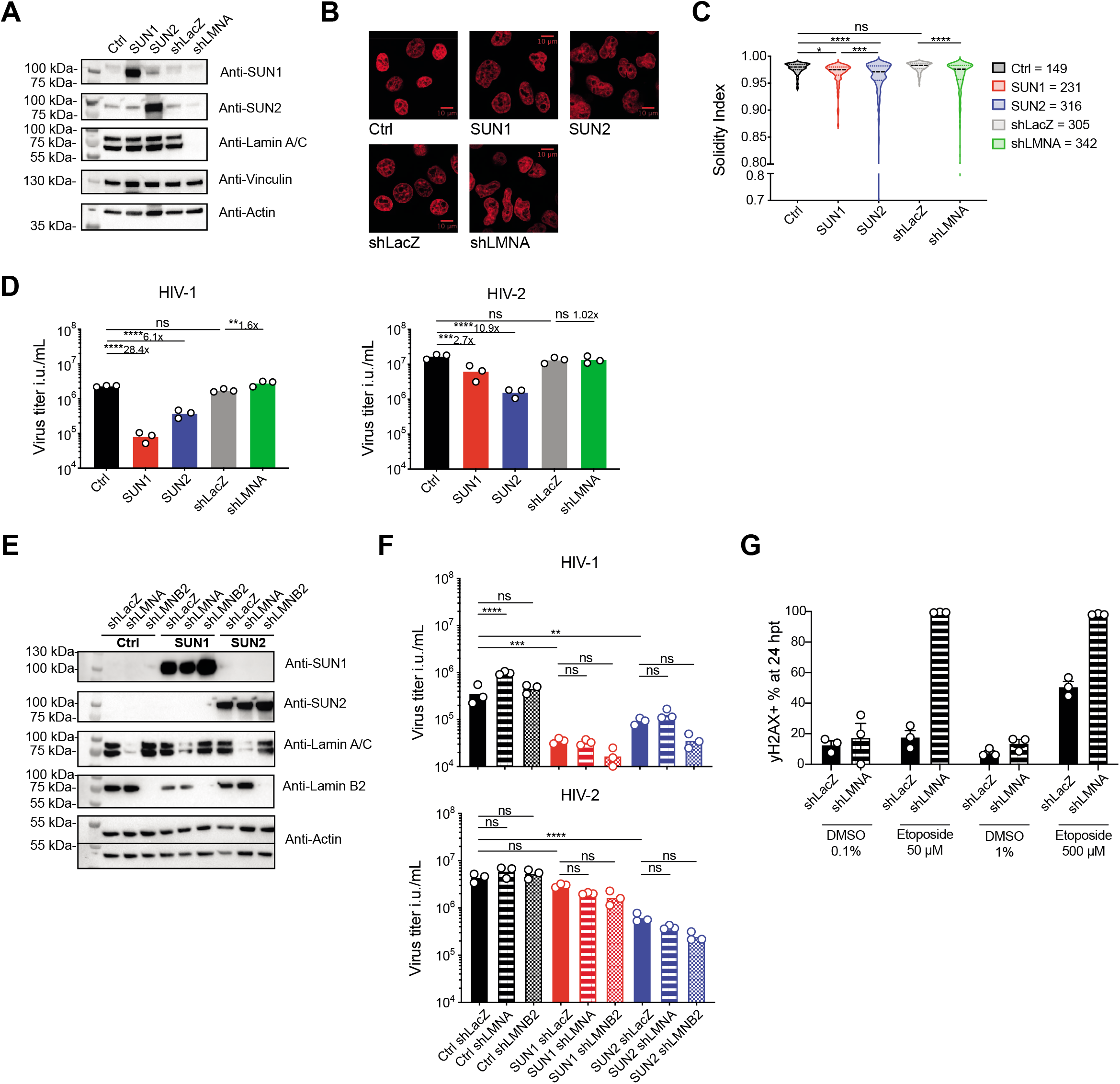
Interplay between HIV-1 infection, Lamin A/C protein and the DNA damage response. **(A)** Detection of SUN1, SUN2, Lamin A/C and actin in HeLa cells transduced with mTagBFP-2A control, SUN1 or SUN2 lentivectors, negative control LacZ (shLACZ) or lamin A/C (shLMNA) targeting shRNA-encoding lentivectors. **(B)** Nuclei of HeLa cells lines as in (A) visualized on fixed cells using SiR-DNA dye. Images show signal from an individual, central confocal plane, scale bar is at 10 μm. **(C)** Solidity index of nuclei as in **(B)**. Legend indicates total number of nuclei analyzed per cell line (representative of n=2 experiments, one on fixed cells and one with live imaging). Unpaired ANOVA one-way with Sidak’s, line at median. **(D)** Viral titers based on percentages of GFP^+^ HeLa cells lines from **(A)** after infection with serial dilutions of HIV-1 and HIV-2 (n=3, paired RM ANOVA one-way with Sidak’s post-test, line at mean). **(E)** Detection of SUN1, SUN2, Lamin A/C, Lamin B2 and actin in HeLa cells co-transduced with mTagBFP-2A control, SUN1 or SUN2 lentivectors and with LacZ, lamin A/C or lamin B2 (shLMNB2) targeting shRNA-encoding lentivectors. **(F)** Viral titers based on percentages of GFP^+^ HeLa cells as shown in **(E)** after infection with serial dilutions of HIV-1 and HIV-2 (n=3, paired RM ANOVA one-way with Sidak’s post-test, line at mean). **(G)** Quantification of gH2AX^+^ HeLa cells transduced with either a LacZ or lamin A/C targeting shRNA lentivector, 24 hours after treatment with indicated doses of etoposide or DMSO control (n=3, line at mean ± SEM). Ctrl = control, hpt = hours post treatment, *p < 0.05, **p < 0.001, ***p < 0.001, ****p < 0.0001; ns, not statistically significant.

Next, we examined the level of γH2AX after etoposide treatment and Lamin A/C depletion. Treatment with a high dose of etoposide (500 μM) for 24 hours induced an increase of γH2AX level in wild-type HeLa cells, while a lower dose (50 μM) had no impact at this time point (**Figure 6G**). In the absence of Lamin A/C, HeLa cells became hyper-sensitive to etoposide treatment (**Figure 6G**). Thus, the increase in HIV-1 infection observed after Lamin A/C depletion correlates with an increased sensitivity to DNA damage. Altogether, these results establish a functional correlation between the effect of SUN protein and endogenous Lamin A/C on HIV-1 infection and on the cellular response to DNA damage induced by an exogenous compound.

### Elevated SUN proteins do not alter NPC density, passive import or cell stiffness

We next sought to identify the mechanisms that delineate the effects of elevated SUN proteins and endogenous Lamin A/C depletion on infection. We characterized biophysical and structural parameters in SUN-expressing cells. HIV-1 enters the nucleus through NPCs. We labelled NPCs using a marker of Nup153 on tangential confocal microscopy sections of the nuclear envelope (**Figure S2A**). Overexpression of SUN proteins did not alter NPC density at the nuclear envelope (**Figure S2B**). To determine if the NPC functionality was impaired by SUN protein overexpression, we measured passive diffusion through the NPC using a Fluorescence Recovery After Photobleaching (FRAP) assay on ubiquitous GFP. SUN proteins had no impact on the rate of recovery of nuclear GFP (**Figure S2C, Movie 1**). Lamin A/C depletion reduces stiffness of the nuclear envelope, resulting in a more deformable nucleus (Lammerding et al., 2004). To determine if SUN proteins expression induced the opposite to match the effects on infection, we measured the viscoelastic properties of the nuclei using a microfluidic micropipette assay (Davidson et al., 2019). While we confirmed that Lamin A/C depleted cells are more deformable, expression of SUN proteins had no impact on nuclear deformability (**Figure S2D**).

### HIV-1 infection requires movement of the chromatin

Next, we turned our attention to the endogenous state of chromatin. We performed live-imaging of cells with a DNA stain after SUN overexpression or Lamin A/C depletion. We observed that SUN1 and SUN2 overexpression appeared to lock the nucleus in place, while nuclei of Lamin A/C-depleted cells appeared highly dynamic (**Figure 7A, Movies 2–6**). We first asked whether the extent of chromatin movement inside the nuclei was altered. We isolated movies of single nuclei and performed a registration step to normalize X-Y positions and angle, therefore suppressing general nuclei displacement and rotation. We next measured chromatin movement in the registered nuclei by performing a particle image velocimetry (PIV) analysis. Strikingly, SUN1 and SUN2 overexpression reduced the displacement of chromatin over time and this effect was more pronounced with SUN1, while Lamin A/C depletion had the converse effect (**Figure 7B, S3A**). HeLa cells also exhibit seemingly random rotation of their nuclei, at various speeds and frequencies. Using the same dataset, we measured the rotation of the whole nucleus relative to the cytoplasm. We corrected the translational displacement of nuclei by registration and measured the angle of rotation over time using a custom-made analysis script (**Movie 7**). SUN1 and SUN2 reduced the average speed of nuclear rotation and the fraction of time spent rotating above a threshold of 1° (**Figure 7C, S3B**). The rotation of Lamin A/C-depleted nuclei was visibly higher than controls, but could not be reliably quantified due to the high levels of chromatin displacement that hampered the ability to set reference points. Overall, these results indicate that the impact of SUN and Lamin A/C proteins on HIV-1 infection is associated with the movement of chromatin within the cells.

**Figure 7.**
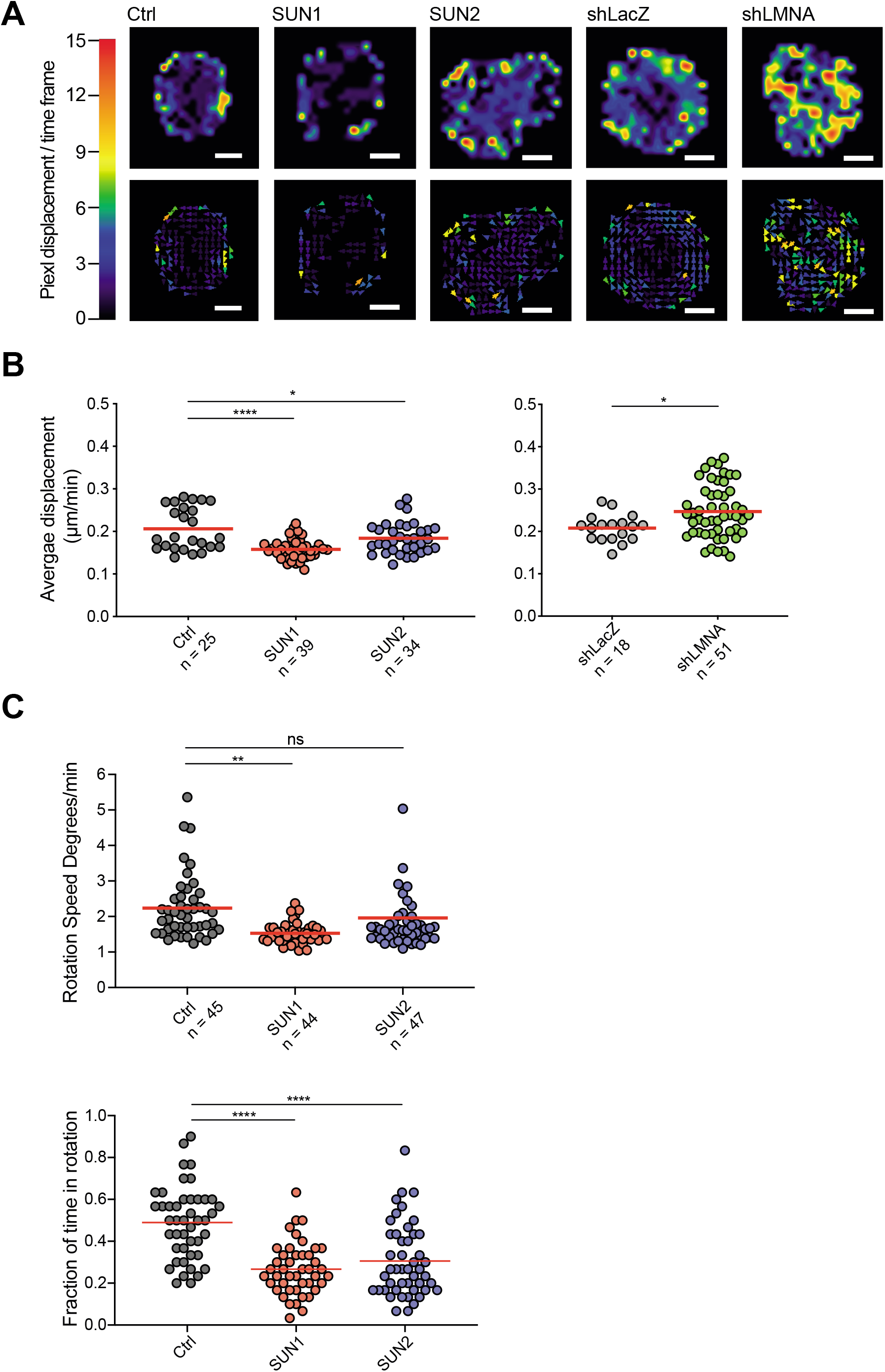
SUN protein overexpression and endogenous Lamin A/C limit movements of the chromatin. **(A)** Particle Image Velocimetry (PIV)–of DNA within nuclei of HeLa cells transduced with mTagBFP-2A control, SUN1 or SUN2 lentivectors, negative control LacZ or lamin A/C targeting shRNA-encoding lentivectors. PIV is shown for individual representative nuclei of each cell line, top panels show overall flow and bottom panels show individual vectorial displacements between two consecutive frames corresponding to two minutes of imaging. Scale bar corresponds to 5 μm. Reference color scale for pixel displacement per time frame is shown on left. **(B)** Quantification of DNA displacement as μm/min from images as in **(A)**. Results are shown for one experiment from n = 2. Left, unpaired ANOVA one-way with Sidak’s post-test, line at mean. Right, unpaired student t-test, line at mean. **(C)** Quantification of nuclear rotation speed as degrees/minute (top) and rotation duration as fraction of total time spent rotating over a threshold of 1° (bottom) in HeLa cells transduced with mTagBFP-2A control, SUN1 or SUN2 lentivectors and imaged as in **(A)**. Results are shown for one experiment from n = 2. Unpaired ANOVA one-way with Sidak’s (top) or Turkey’s (bottom) post-test, line at mean. Ctrl = control, *p < 0.05, **p < 0.01, ****p < 0.0001; ns, not statistically significant.

## Discussion

Our findings reveal that SUN1 and SUN2, though paralogs, have distinct effects on HIV infection. SUN1 overexpression is more efficient at inhibiting HIV-1 than SUN2. Meanwhile, SUN2 overexpression shows a marked antiviral activity against HIV-2. An analysis of the viral step impacted by these two proteins also reveals a discrepancy: while both SUN1 and SUN2 overexpression reduces the level of integrated HIV-1 DNA, SUN1 also inhibits total viral DNA amount while SUN2 reduces the levels of 2-LTR circles, a hallmark of nuclear entry. We also reveal that SUN1 and SUN2 differ in their response to DNA damage and its impact on HIV infection. SUN1 limits the response to etoposide as measured by the levels of γH2AX, while SUN2 enhances it. Strikingly, etoposide largely rescues the antiviral effect of SUN2 overexpression on HIV-1, while SUN1 is resistant to this effect. This result suggests that SUN1 and SUN2 may differ in the ways in which they establish interactions and functions within the nucleus, in line with previously reports showing non-redundant effects of SUN1 and SUN2 (Lei et al., 2009; Liu et al., 2007; Zhu et al., 2017)

Etoposide treatment also enhances the infection by HIV-1 by two-fold in HeLa cells in the absence of SUN protein overexpression, while HIV-2 is largely unaffected. Such a proviral effect of DNA damage has been previously observed in conditions of integrase inhibition (Ebina et al., 2012; Koyama et al., 2013). In contrast, it was previously reported that etoposide treatment inhibits HIV-1 infection in monocyte-derived macrophages (Mlcochova et al., 2018). We speculate that this is explained by the presence of a SAMHD1-dependent block in this cell type induced by etoposide in macrophages, but not in HeLa cells.

Similar to etoposide, ATR inhibition also enhances HIV-1 infection by 2-fold. The lack of requirement for ATR in HIV infection is consistent with prior studies (Ariumi et al., 2005; DeHart et al., 2005). Interestingly, the effect of ATR inhibition requires the presence of the Vpr gene in HIV-1. As a virus-encoded gene, Vpr has be shown to induce an ATR-dependent G2 arrest of the cell cycle (Zimmerman et al., 2006). Using purified and p24-normalized virus preparations, we made the unexpected observation that the Vpr-deficient virus is actually as infectious as the wild-type virus stimulated with ATR inhibition. In other words, in this asynchronous system of single-round infection of HeLa cells, the presence of the Vpr gene appears to provide a counter-intuitive two-fold reduction in infectivity of the virus, which is alleviated by ATR inhibition. However, Vpr has been associated with an enhanced expression for the viral LTR during G2 arrest (Goh et al., 1998). We speculate that in terms of viral replicative fitness, the reduction in single-round infectivity entailed by Vpr is cancelled out by this proviral effect of Vpr during G2 arrest. Of note, both SUN1 and SUN2 overexpression inhibit HIV-1 infection irrespective of ATR inhibition. In contrast, etoposide rescues the antiviral effect of SUN2 on HIV-1, raising the possibility that ATR itself might play a role in the rescue of the SUN2 antiviral effect.

Multiple lines of evidence from our work indicate that the structure of the nuclear envelope impacts HIV infection. The ability of SR-KASH to partially rescue the antiviral effects of SUN proteins indicates that the LINC complex, which is located at the nuclear envelope, is involved. Despite the important morphological changes induced by SUN protein expression, we did not observe any change at the level of gene expression, suggesting that any SUN-mediated effect on the nucleus and, subsequently, on HIV infection is post-transcriptional. ATR is enriched at the nuclear envelope during S phase and upon mechanical stretching, two processes that increase nuclear envelope stress (Kumar et al., 2014). ATR-deficient cells exhibit deformed nuclei, reminiscent of Lamin A/C depletion or SUN protein overexpression (Kidiyoor et al., 2020). Vpr overexpression was previously shown to induce herniations of the nuclear envelope associated with defects in the nuclear lamina (de Noronha et al., 2001). We also find that endogenous Lamin A/C has an antiviral effect, in agreement with a previous study (Sun et al., 2018).

We examined several effects of SUN protein overexpression and endogenous Lamin A/C on nuclear shape, deformability, NPC distribution and function and chromatin dynamics. The effects on HIV-1 infection match the effects of the proteins on chromatin dynamics: decreased HIV-1 infection is associated with a decreased chromatin motility inside the nucleus and with decreased rotation of the nucleus relative to the cytoplasm. In contrast, HIV-2 infection is more susceptible to SUN2 than SUN1 and is not affected by Lamin A/C depletion. Overexpression of SUN2 deforms nuclei more than SUN1 but internal chromatin dynamics and nuclear rotation are less impacted. This strain specificity could be linked to a different dependency on host factors between HIV-1 and HIV-2 (Braaten and Luban, 2001). SUN and Lamin proteins have been previously linked to chromatin mobility and nuclear rotation (Ji et al., 2007; Lottersberger et al., 2015; Oza et al., 2009; Ranade et al., 2019). Interestingly, nuclear rotation is required for optimal infection by another nuclear-invading virus, HCMV (Procter et al., 2018). This rotation is required to promote spatial chromatin segregation that favors viral gene expression (Procter et al., 2018). We propose that HIV-1 infection requires nuclear rotation and chromatin movements for optimal integration and subsequent viral expression.

Our results highlight the interplay between HIV infection, structural proteins of the nuclear envelope and the DNA damage response. Nuclear rotation and chromatin dynamics emerge as potentially important factors that control HIV infection. Future studies will be required to address the underlying molecular mechanisms, which we anticipate will require the use of biophysical approaches. It will also be important to examine these mechanisms in the frame of the diversity of lentiviruses and their relevance for viral replication and innate immune sensing mechanisms in primary target cells.

## Acknowledgments

We thank N. De Silva for setting up the gH2AX intranuclear staining; V. Teixeira Rodrigues for assistance with primary macrophages; P. Benaroch for critically reading this manuscript; M. Piel, N. De Silva, N. Jeremiah and A. Williart for discussions. We acknowledge the PICT-IBiSA imaging facility, member of the France-BioImaging national research infrastructure, supported by the CelTisPhyBio Labex (ANR-10-LBX-0038) part of the IDEX PSL (ANR-10-IDEX-0001-02 PSL), and Audrey Rapinat and David Gentien from the Genomics Platform at Institut Curie. This work was supported by Institut Curie, INSERM, and by grants from the Agence Nationale de la Recherche (ANR-10-IDEX-0001-02 PSL, ANR-11-LABX-0043, ANR-17-CE15-0025-01, ANR-19-CE15-0018-01, ANR-18-CE92-0022-01, France-BioImaging ANR-10-INSB-04), the Agence Nationale de la Recherche sur le SIDA (ECTZ36691, ECTZ25472, ECTZ71745), Sidaction (VIH2016126002, 17-1-AAE-11097-2). AB was supported by fellowships from PSL University and Fondation pour la Recherche Médicale, grant number 8250. PMD was supported by fellowships from La Ligue contre le Cancer (REMX17751) and Fondation ARC (PDF20161205227).

## Supplemental Material

**Figure S1.**
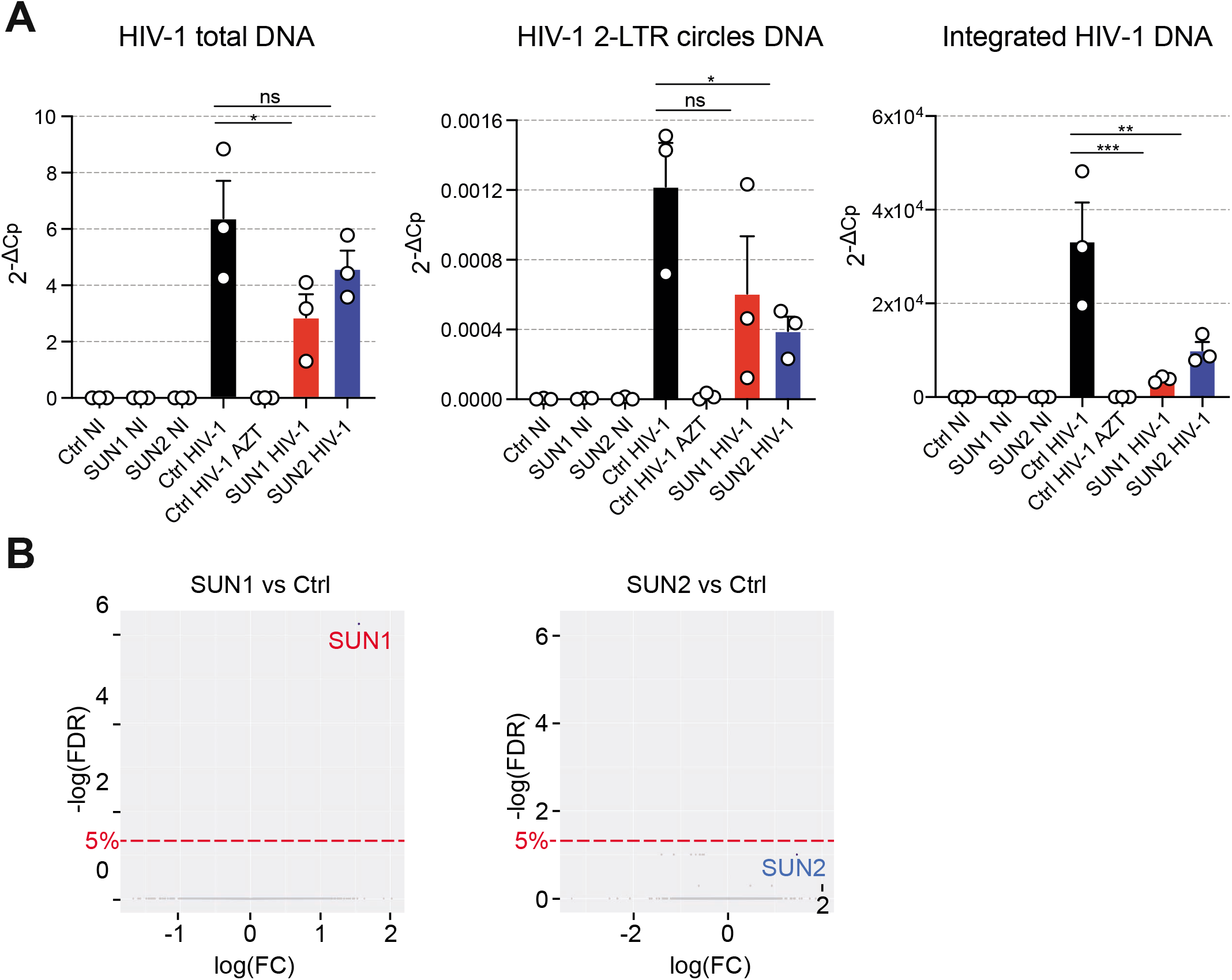
HIV-1 DNA quantification and gene expression analysis of SUN-overexpressing cells. **(A)** Detection of HIV-1 total DNA, 2-LTR circles DNA and integrated DNA by RT-qPCR at 24 hours after infection (dilution factor: 0.1) of HeLa cells transduced with mTagBFP-2A control, SUN1 or SUN2 lentivectors. AZT (24 μM) was added during infection when indicated (n=3, paired RM ANOVA one-way with Turkey’s post-test, line at mean ± SEM). **(B)** Differential gene expression in SUN1 (Left) and SUN2 (Right) overexpressing cells over control cells (FC, fold change; FDR, false discovery rate). Overexpressed SUN2 was codon-optimized, rendering it sub-optimal for probe-based detection. Ctrl = control, *p < 0.05, **p < 0.001, ***p < 0.001; ns, not statistically significant.

**Figure S2.**
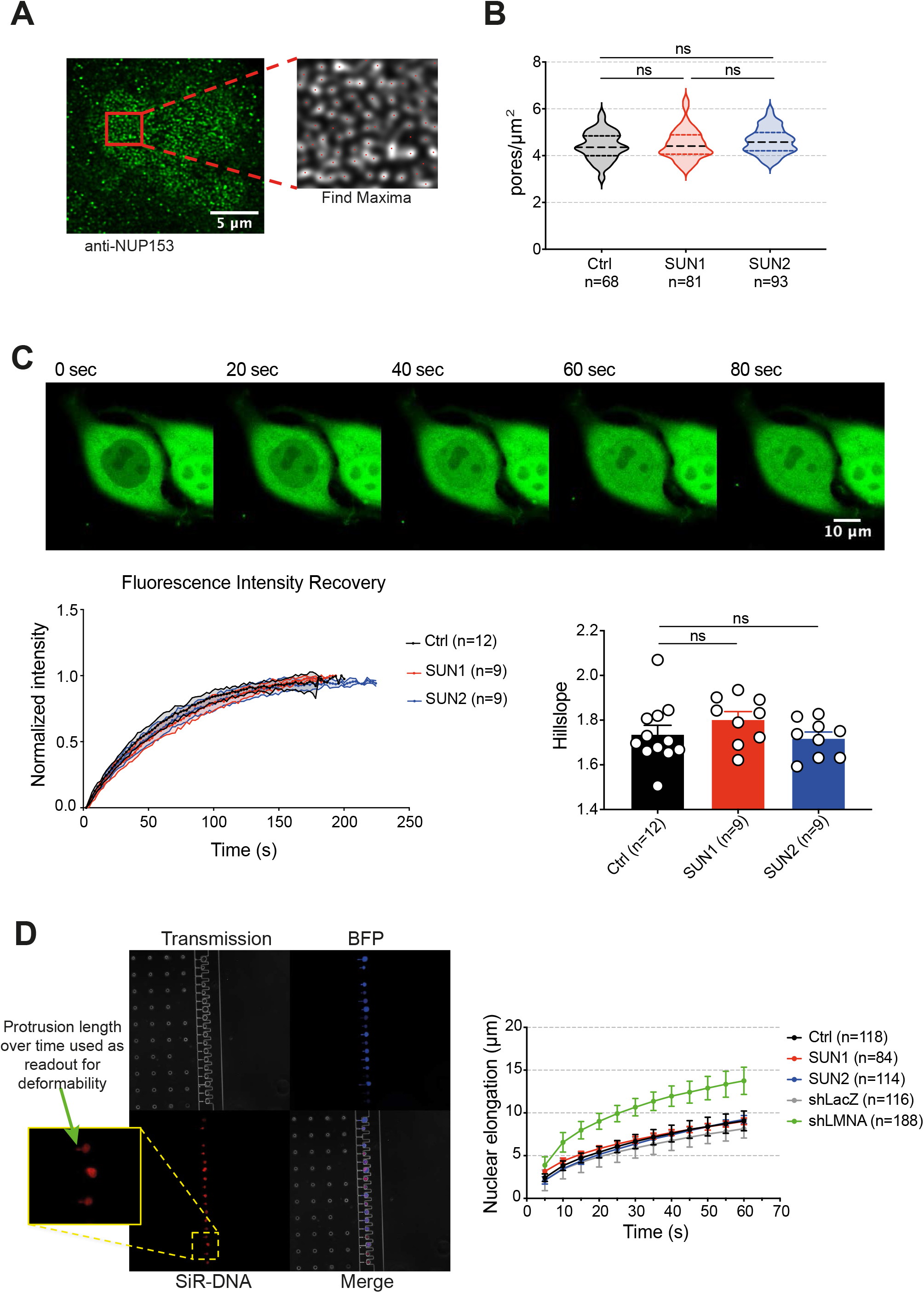
Analysis of NPC density, passive nuclear import and cellular stiffness of SUN-overexpressing cells. **(A)** Left, confocal imaging of NUP153 staining in HeLa cells. Right, blow-up or region of interest and detection of individual of NPC (red) using “Find Maxima” function. A representative nucleus of cells overexpressing mTagBFP-2A-SUN2 is shown. **(B)** Quantification of NPC density per μm^2^ in nuclei from HeLa cells transduced with mTagBFP-2A control, SUN1 or SUN2 lentivectors, imaged and analyzed as in **(A)** (total number of nuclei analyzed per cell line are indicted, n=1 experiment; un-paired ANOVA one-way with Sidak’s post-test, line at mean). **(C)** Passive diffusion from the cytoplasm to the nucleus measured by FRAP, in HeLa cells transduced with mTagBFP-2A control, SUN1 or SUN2 lentivectors and a lentivector encoding GFP. Top, representative layout of a photobleached control HeLa cell: imaging starts at t = 0 after high intensity laser exposure and ends at t = 80 seconds, when fluorescence in the nucleus has been recovered (scale bar: 5 μm). Bottom left: GFP intensity recovery over time (s). For each individual cell, the background intensity was subtracted and intensity was normalized to 0 at t0 after photobleaching while the max intensity reached during the course of each measurement was set to 1. Curves show mean and standard deviation per time point across indicated number of cells per condition. Bottom right, hillslopes for each cell were calculated via non-linear regression fit (total number of nuclei analyzed per cell line are indicated, n=1 experiment; un-paired ANOVA one-way with Dunnett’s post-test, line at mean ± SEM). **(D)** Measurement of nuclear deformability. Left, representative image showing tagBFP-2A-Ctrl HeLa cells stained with SiR-DNA and going through microchannels under externally-applied pressure. The green arrow indicates the elongation of SiR-DNA staining within the channel, which is measured over time as a readout for nuclear deformability. Right, quantification of nuclear deformability across HeLa cell lines described in **(6A)** over time (the number of nuclei measured per cell line is indicated within brackets, n = 3 independent experiments). Ctrl = control, ns, not statistically significant.

**Figure S3.**
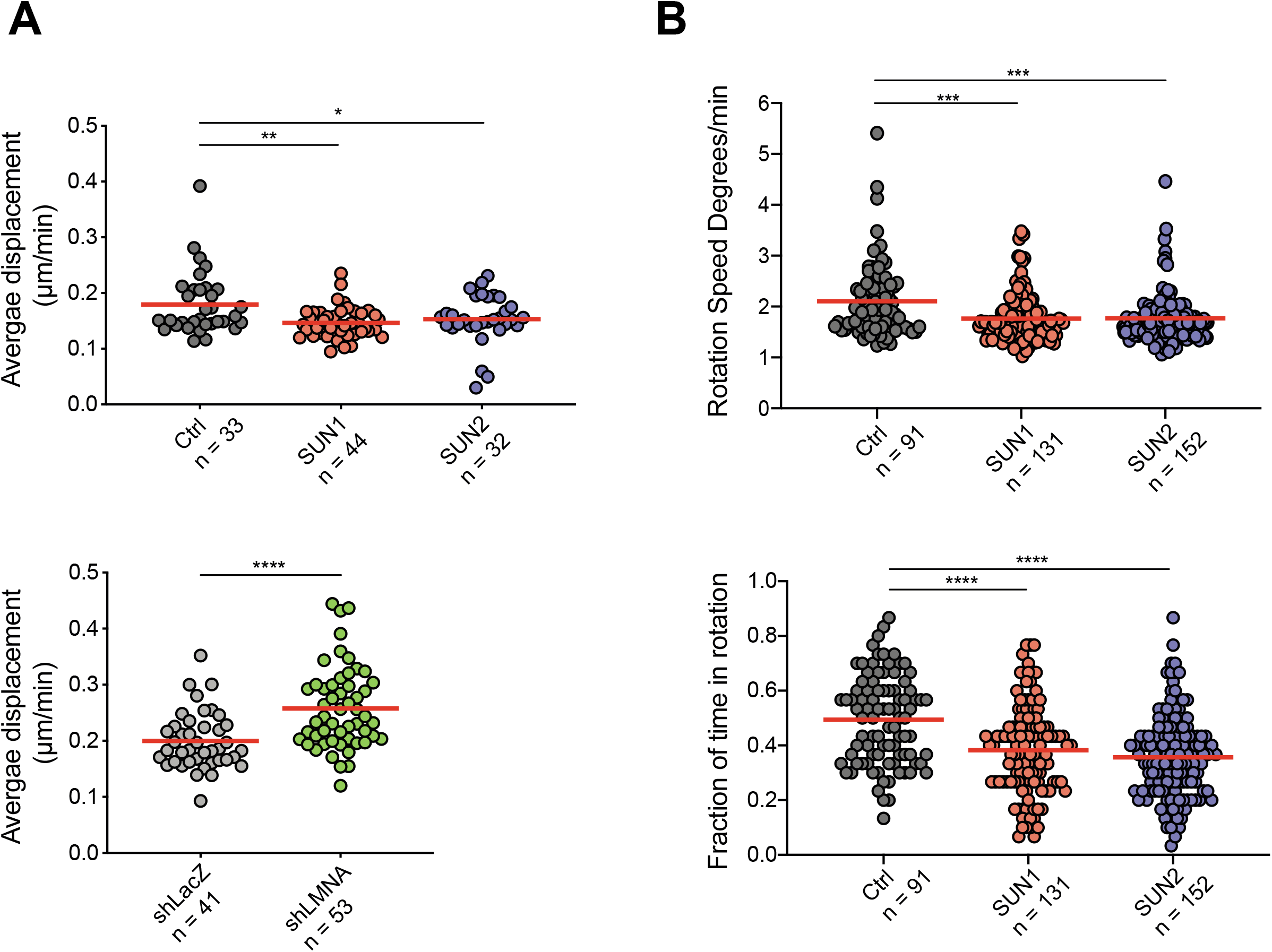
SUN protein overexpression and endogenous Lamin A/C limit movements of the chromatin, second experiment. **(A)** Quantification of DNA displacement as μm/min from images as in Figure 7B, second experiment. Top, unpaired ANOVA one-way with Sidak’s post-test, line at mean. Bottom, unpaired student t-test, line at mean. **(B)** Quantification of nuclear rotation speed as degrees/minute (top) and rotation duration as fraction of total time (bottom) spent rotating over a threshold of 1°, as in Figure 7C, second experiment. Unpaired ANOVA one-way with Sidak’s (top) or Turkey’s (bottom) post-test, line at mean. Ctrl = control, *p < 0.05, *p < 0.01 ***p < 0.001, ****p < 0.0001; ns, not statistically significant.

**Movie 1 Passive diffusion from the cytoplasm to the nucleus measured by FRAP.**

Representative movie of a HeLa cell overexpressing mTagBFP-2A-Ctrl and ubiquitous GFP lentivectors. Immediately prior to imaging, high intensity 488 nm laser was directed to a small region within the nucleus to bleach the GFP signal within this compartment. Imaging of the whole cell starts at t = 0 immediately after bleaching. Recovery of the GFP signal in the nucleus is observed as GFP diffuses back in from the cytoplasm until equilibrium is obtained. One frame was taken every 2 seconds.

**Movies 2 Quantification of chromatin dynamics in mTagBFP-2A-overexpressing control HeLa cells.**

Chromatin dynamics assessed by PIV. The analysis was performed on live confocal imaging of DNA staining using SiR-DNA. An image was taken every 2 minutes. The left panel shows registration of the object (one nucleus) across 10 time frames while the right panel shows the flow of pixel clusters (interrogation windows) within the object, across consecutive time frames. Warmer colors indicate high levels of displacement while cold colors indicate more static regions. Scale bar corresponds to 5 μm.

**Movie 3 Quantification of chromatin dynamics in mTagBFP-2A-SUN1-overexpressing HeLa cells.**

Analysis was performed as for Movie 2.

**Movie 4 Quantification of chromatin dynamics in mTagBFP-2A-SUN2-overexpressing HeLa cells.**

Analysis was performed as for Movie 2.

**Movie 5 Quantification of chromatin dynamics in HeLa cells expressing a control shRNA against LacZ.**

Analysis was performed as for Movie 2.

**Movie 6 Quantification of chromatin dynamics in HeLa cells expressing a shRNA against Lamin A/C.**

Analysis was performed as for Movie 2.

**Movie 7 Measurement of nuclear rotation in a HeLa cell.**

The analysis was performed on live confocal imaging of DNA staining using SiR-DNA. An image was taken every 2 minutes, for a total duration of 60 minutes. The angle was measured in reference to an image containing 2 fixed points (one at the center and one on the edge of the nucleus). In this example, HeLa overexpressing mTagBFP-2A control are shown.

## Materials and methods

### Constructs

The plasmid constructs for lentiviral expression and HIV infection used in this study are listed in **Table 1**. pTRIP-SFFV-tagBFP-2A-SUN1 Dharmacon was generated by overlapping PCR cloning from commercially bought cDNA (MGC cDNA cloneID: 40148817) into pTRIP-SFFV-tagBFP-2A (Cerboni et al., 2017). pTRIP-SFFV-tagBFP-2A-ntSUN2 was generated by overlapping PCR mutagenesis from pLX304-SUN2 (Lahaye et al., 2016) into pTRIP-SFFV-tagBFP-2A with concomitant introduction of silent mutations that are not targeted by SUN2 shRNA 4 and 5 (respectively GAGCCTATTCAGACGTTTCACTTT to GAACCGATCCAAACTTTCCATTTC and AAGAGGAAATCCAGCAACATGAAG to AAACGCAAGAGTTCTAATATGAAA). pTRIP-SFFV-tagBFP-2A-SUN1 Dharmacon (1-298)-ntSUN2 (220-717) and pTRIP-SFFV-tagBFP-2A-ntSUN2 (1-219)-SUN1 Dharmacon (299-785) were generated by overlapping PCR cloning from the full-length constructs. pTRIP-SFFV-tagRFP657-2A-SUN1 and pTRIP-SFFV-tagRFP657-2A-ntSUN2 were generated via restriction enzyme digestion from the tagBFP expressing vectors and ligation into pTRIP-SFFV-TagRFP657-2A backbone. HIV-GFP env-nef- was generated by PCR-mediated insertion of the Vpr+Vif+Vpu+ cassette from NL4-3 into HIV-GFP (Manel et al., 2010). HIV-GFP env-nef-vpr- was generated by overlapping PCR mutagenesis from HIV-GFP env-nef-, introducing a frameshift mutation within *vpr*, after the codon corresponding to amino-acid I63 (gaattc to gaaTTAAttc). HIV-mTagBFP2 and HIV-2 ROD9 ΔenvΔnef mTagBFP2+ were obtained via overlapping PCR mutagenesis, replacing GFP with the mTagBFP from pTRIP-SFFV-mTagBFP-2A.

### Cells

GHOST (GHOST X4R5), 293FT and HeLa cells were cultured in DMEM with Glutamax, 10% fetal bovine serum (FBS) (Corning), and penicillin-streptomycin (Gibco). Human peripheral blood mononuclear cells (PBMCs) were isolated from buffy coats from normal human donors (approved by the Institut National de la Santé et de la Recherche Médicale ethics committee) using Ficoll-Paque PLUS (GE). CD14^+^ cells were isolated by a positive selection with anti-human CD14 magnetic beads (Miltenyi) from PBMCs. To obtain macrophages (MDMs), CD14^+^ cells were cultured in RPMI with Glutamax, 5% FBS (Eurobio), 5% human serum (Sigma), Penicillin-Streptomycin, Gentamicin (50 μg/ml, GIBCO) and HEPES (GIBCO) in the presence of recombinant human M-CSF (Miltenyi) at 50 ng/ml. Fresh media was added at day 5 or 6, and cells were treated/infected at day 9, after detachment via incubation with StemPro Accutase Cell Dissociation Reagent (Gibco) for 30 minutes at 37°C. Drug treatments performed on cultured cells are listed in **Table 2**.

### Virus production

Viral particles were produced by transfection of 293FT cells in 6-well plates with 3 μg DNA and 8 μl TransIT-293 Transfection Reagent (Mirus Bio) per well. For VSV-G pseudotyped SIVmac virus-like particles (VLPs), 0.4 μg CMV-VSVG and 2.6 μg pSIV3^+^ was used. For VSV-G pseudotyped HIV-1 and HIV-2 GFP or BFP-reporter viruses, 0.4 μg CMV-VSVG and 2.6 μg HIV DNA was used. For overexpression or sh-RNA mediated knock-down, 0.4 μg CMV-VSVG, 1 μg psPAX2 and 1.6 μg of lentivector of interest were combined. One day after transfection, media was removed, cells were washed once, and 3 ml per well of RPMI medium with Glutamax, 10% FBS (Gibco), PenStrep (Gibco), 50μg/ml Gentamicin (Gibco) and 0.01 M HEPES (Gibco) were added. Viral supernatants were harvested 1 day later, filtered using 0.45 μm pore filters, used fresh or aliquoted and frozen at −80°C. When required, the virus was purified and concentrated on a 20% sucrose cushion in phosphate buffered saline (PBS) in Ultra Clear Centrifuge tubes (Beckman Coulter), via ultracentrifugation at 4°C at 31,000 *x g* in a SW32Ti swinging bucket rotor (Beckman Coulter). Viral pellets were then resuspended in complete medium at a 100-fold concentration compared to crude. Viral titers were measured on GHOST cells (titration as previously described (Manel et al., 2010) or using HIV-1 p24 ELISA (XpressBio). ELISA absorbance acquisitions were acquired on a FLUOstar OPTIMA (BMG Labtech) and data were analyzed and exported to Excel with MARS Data Analysis Software (BMG Labtech).

### Cell Transduction for protein overexpression or knockdown

HeLa cells were counted and seeded in 6-well plates on the day prior to transduction. Purified virus was added at a 2:1 volume ratio on medium containing protamine at a final concentration of 1 μg/ml. CD14^+^ monocytes were seeded in 10-cm dishes in the presence of 50 ng/ml M-CSF to induce differentiation into macrophages and transduced with purified SIVmac VLPs and lentiviruses carrying vector of interest, mixed at a 1:1 ratio. Human serum was added at day 1 post transduction. Transductions of monocytes was performed in the presence of protamine at a final concentration of 1 μg/ml. HeLa cells were washed once in PBS and passaged at 48 hours post transduction with or without 2 μg/ml of puromycin. For MDMs, medium was replaced at day 5-6 post transduction. Overexpression was assessed by quantification of fluorescent reporter signal via flow cytometry on a BD FACSVerse flow cytometer. Both overexpression and protein knock-down were confirmed by Western Blotting at day of experiment.

### Cell infection

HeLa, GHOST and MDMs (day 8-9 post transduction) were seeded and infected in the presence of 1 μg/ml of protamine with serial dilutions of frozen viral stocks in a BSL-3 laboratory. Virus was removed at 48 hours post-infection, cells were washed, harvested, stained for viability using Fixable Viability Dye eFluor 780 in PBS where required, fixed in 1% paraformaldehyde (PFA; Electron Microscopy Sciences) and analyzed for GFP or BFP positivity via flow cytometry on a BD FACSVerse flow cytometer. Viral titers were calculated based on seeded cell number and the percentages of infected cells, within the linear range of infection.

### HIV DNA quantification

HeLa cells and MDMs were infected as described, with the addition of infected wells treated with RT inhibitors as negative control. For this purpose, either 24 μM of AZT (Sigma) or 10 μM of NVP (Sigma) were used. After 24 hours, cells were washed in PBS and harvested. Total DNA was extracted from cell pellets using NucleoSpin Tissue (Macherey-Nagel) kit, as per manufacturer’s protocol. Real-time PCR analysis was performed as previously described (Lahaye et al., 2013). Each sample was measured in triplicate for all primers. For beta-globin, primers were bglobin-f and bglobin-r. Cycling conditions were 1x 95°C for 5’; 35x 95°C for 10’’, 65°C for 20’’ (50°C for beta-globin) and 72°C for 30’’. Relative concentrations of total DNA (Late RT), 2-LTR circles and integrated viral DNA were calculated relative to beta-globin using the ΔCt method. The primers used are listed in **Table 3**.

### Western Blotting

0.5 to 1 million cells were lysed in 100 μL of RIPA buffer (50mM Tris HCl, 150mM NaCl, 0.1% SDS, 0.5% DOC, 1% NP-40, Protease inhibitor (Roche; 1187358001)). Lysis was performed on ice for 30’. Lysates were cleared by centrifugation at 8000 g for 8 minutes at 4°C, 20 μl of Laemmli 6x (12% SDS, 30% Glycerol, 0.375M Tris-HCl pH 6.8, 30% 2-mercaptoethanol, 1% bromophenol blue) was added and samples were boiled at 95°C for 15’. Cellular protein lysates were resolved on Criterion or 4%–20% Bio-Rad precast SDS-PAGE gels and transferred to PVDF membranes (Bio-Rad). Membranes were saturated and proteins were blotted with antibodies in 5% non-fat dry milk, PBS 0.1% Tween buffer. ECL signal generated via Clarity Western ECL substrate (Bio-Rad) was recorded on the ChemiDoc-XRS or ChemiDoc Touch Bio-Rad Imager. Data was analyzed and quantified with the Image Lab software (Bio-Rad). The antibodies used in this study are listed in **Table 4**.

### Live Confocal Imaging

For live imaging, HeLa cells were plated either in a glass bottom FluoroDish (World Precision Instruments) or in a glass-bottom Cellview Cell Culture Dish with 4 compartments (Greiner Bio-One), on the day prior to experiments. One hour before imaging, cells were incubated with 1 μM of SiR-DNA (Tebu Bio), directly in the culture medium, at 37°C. Images of cells were acquired with a Leica DmI8 inverted microscope equipped with an SP8 confocal unit using a 20x dry objective (NA=0.75, pixel size was fixed to 0.284 μm). Imaging was performed in an on-stage incubator chamber at 37°C, with 5% CO_2_. An image per condition was taken every 2 minutes, unless specified otherwise.

Image analysis was performed using Fiji software (Schindelin et al., 2012). For chromatin dynamics analysis, a homemade macro was first used to do segmentation of each nucleus on the movie and identify them using the 3D object counter. Particle Image Velocimetry (PIV) analysis was then performed on SiR-DNA staining using the PIV plug-in (Tseng et al., 2012). PIV is a basic optic flow analysis, that divides each image of a stack in small clusters of pixels (interrogation windows) and measures the displacement of each cluster between pairs of consecutive frames. The cross-correlation then generates a pattern of “movements” within the nucleus that are color-coded based on the amplitude of the vector corresponding to the displacement of each cluster. An in-house script was used to first align each individual nucleus, then measure and average SiR-DNA displacements over the ten first time points. Red shades indicate higher amplitudes of displacement while violets correspond to quasi-immobile clusters. For nuclear rotation analysis, a macro was used to measure rotation angles across the first 30 frames. Briefly, individual nuclei were first aligned using a translation transformation of the MultiStackReg plug-in (Brad Busse, Stanford), then they were aligned using the rotation transformation and the transformation was applied to a reference image containing 2 fixed points (one at the center and one on the edge) to measure the rotation. A threshold of 1 degree/minute was used to define rotating nuclei. The percentage of rotation time and the average velocity was then computed.

### Confocal Immunofluorescence Imaging

For immunofluorescence, HeLa cells were grown overnight onto 12 mm glass coverslips (Thermo Scientific) placed in 6-well plates. Cells were fixed with 4% PFA for 20 minutes at room temperature. Coverslips were washed multiple times with PBS and quenched with 0.1M Glycine in PBS(Life Technologies) for 10 minutes at room temperature. Coverslips were then blocked with PBS, 0.2% (w/v) bovine serum albumin (BSA) (Euromedex), 0.05% (w/v) Saponin from quillaja bark (SIGMA) for 30 minutes at room temperature. Cells were stained overnight with anti-NUP153 antibody at 2 μg/mL (1:50 dilution) or with Normal Rabbit IgG Isotype Control at corresponding concentration of the primary antibody, in PBS, 0.2% (w/v) BSA, 0.05% (w/v) Saponin + 10% goat serum (Sigma), at 4°C in a humidified chamber. The following day, cells were washed multiple times and incubated with the secondary antibody Alexa Fluor 546 goat anti-rabbit IgG (H+L) (Invitrogen; 1:200 dilution in PBS-BSA-Saponin) in the presence of 1 μM of SiR-DNA for 2 hours in the dark, at room temperature. Coverslips were washed multiple times in PBS-BSA-Saponin and finally rinsed once in distilled water. Coverslips were mounted onto glass slides using Fluoromount G (eBioscience) mounting medium. The slides were finally dried at 37°C for 1h and stored at 4°C. Cells were imaged with a Leica DmI8 inverted microscope equipped with an SP8 confocal unit using an oil immersion 63x objective (NA=1.4) with applied Type F Immersion Liquid (Leica).

### Fluorescence Recovery After Photobleaching

Cells were seeded at 2.5 x 10^5^ cells/dish in a glass bottom FuoroDish (World Precision Instruments) on the day prior to the experiment. Cells were imaged with a Leica DmI8 inverted microscope equipped with an SP8 confocal unit using a 20x dry objective (NA=0.75). Imaging was performed in an on-stage incubator chamber at 37°C, with 5% CO_2_. Two independent modules were used in a sequential manner: one for bleaching, one for imaging the signal recovery. During the application of the bleaching module, the 488 laser was focused at an intensity of 5% and with a gain of 0.1% on to an area within the nucleus of each cell at maximum zoom for 20 seconds. Immediately afterwards, the first sequence was manually cancelled, the resolution was optimized, imaging area was restored to the whole cell for the second sequence. The laser power was set for optimal imaging level and images of the whole cell were acquired for 3 min circa at the rate of one image every 2.2 seconds.

### Intracellular staining for flow cytometry

Cell surface staining was performed in PBS, 1% BSA (Euromedex), 1mM EDTA (GIBCO), 0.01% NaN3 (AMRESCO) (FACS Buffer) at 4°C. Viability staining (Live-Dead) with Fixable Viability Dye eFluor 780 was performed in PBS at 4°C. Cells were resuspended in FACS Buffer prior to final acquisition. Intracellular staining of gH2AX was performed using the FOXP3/Transcription Factor Staining Buffer Set (eBioscience) as per manufacturer’s protocol. Cells were resuspended in FACS Buffer prior to final acquisition. All flow cytometry acquisitions were performed on the FACSVerse (BD) using the FACSSuite software (BD) and analyzed on FlowJo v10. The antibodies used are listed in **Table 4**.

### Electron Microscopy

Cells were seeded at 5 x 10^4^ cells/well in a 24w plate onto sterile 12 mm glass coverslips (Thermo Scientific) and left to adhere overnight. The following morning, cells were washed in PBS and were fixed using 2% glutaraldehyde in 0.1 M cacodylate buffer, pH 7.4 for 1h, post fixed for 1h with 2% buffered osmium tetroxide, dehydrated in a graded series of ethanol solution, and then embedded in epoxy resin. Images were acquired with a digital 4k CCD camera Quemesa (EMSIS GmbH, Münster, Germany) mounted on a Tecnai Spirit transmission electron microscope (ThermoFisher, Eindhoven, The Netherlands) operated at 80kV.

### Micropipette aspiration microscopy

Prior to harvest, HeLa cell lines were incubated with 1μM SiR-DNA dye from Tebu Bio for 1h30 at 37°C in cell culture medium. Cells were washed, harvested and resuspended at a concentration of 5×10^6^ cells/mL in sterile 3% BSA in PBS-0.2% FBS. Cells were subjected to the experimental conditions as described previously (Davidson et al., 2019).

### Microarray Gene Expression (Affymetrix)

Total RNA was extracted from 10^6^ HeLa cells using NucleoSpin RNA and adjusted to 50 ng/μL. A WT PLUS amplification and labeling protocol was conducted with 100 ng of total RNA. Samples passed the quality control with a high score. The Affymetrix analysis was performed by the NGS platform at Institut Curie using the Human Gene 2.0 ST chip. Human Gene 2.0ST array were scanned using a Genechip 7G scanner, according to the supplier’s protocol. Micro-array analyses were processed with R using packages from Bioconductor. The quality control was performed using ArrayQualityMetrics package without detecting any outlier among the experiment. Data was normalized using the Robust Multi-Array Average algorithm from the Oligo package. Annotation of the probes was done using the hugene20 annotation data (chip hugene20sttranscriptcluster) from Bioconductor. Differential gene-expression analysis was performed with Limma. The accession number for the raw data files is NCBI GEO: GSE162019.

### Statistical Analysis

Statistical analyses were performed in Prism 7 or 8 (GraphPad Software) as indicated in the figure legends.

## Supplementary Tables

**Table 1:**
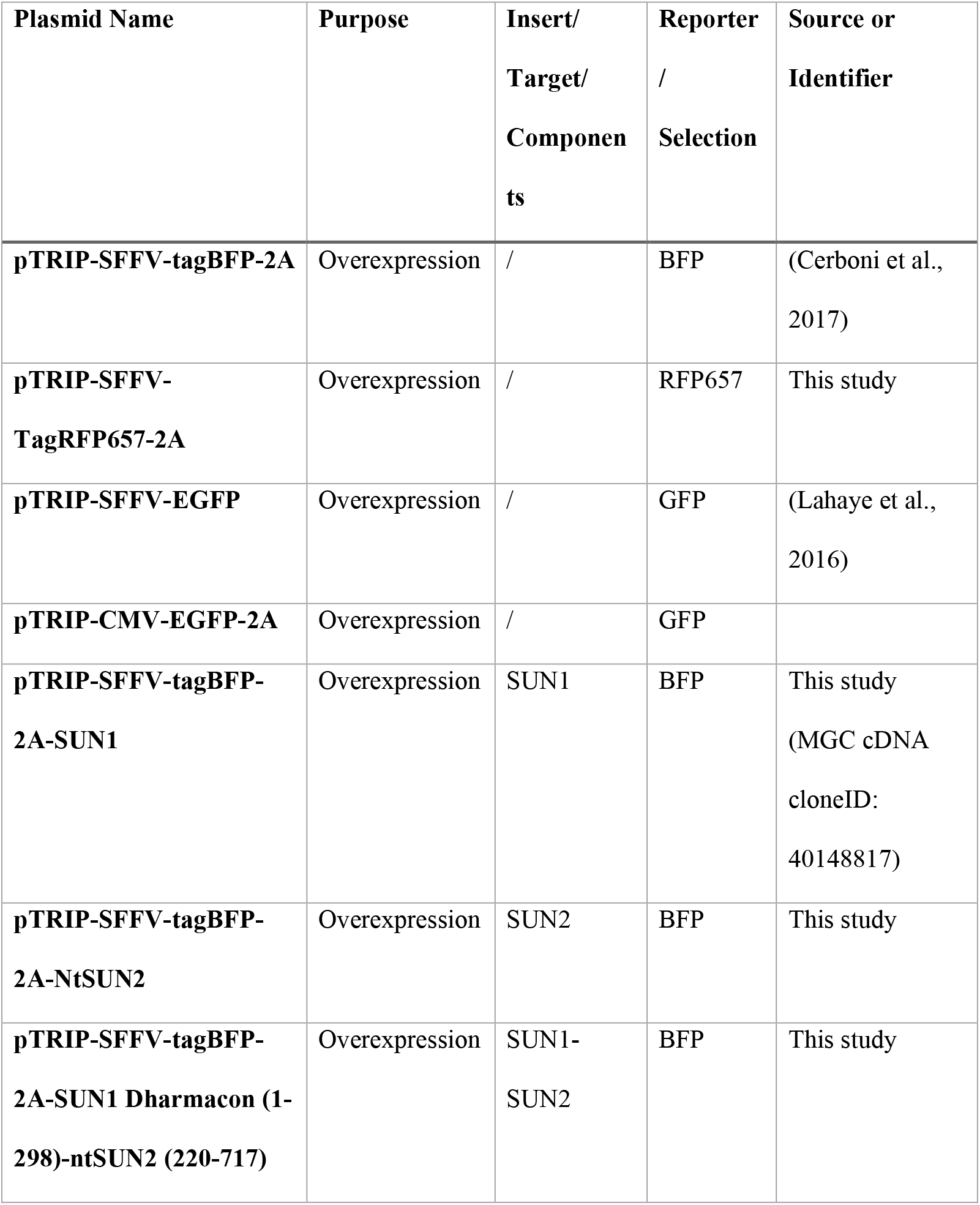
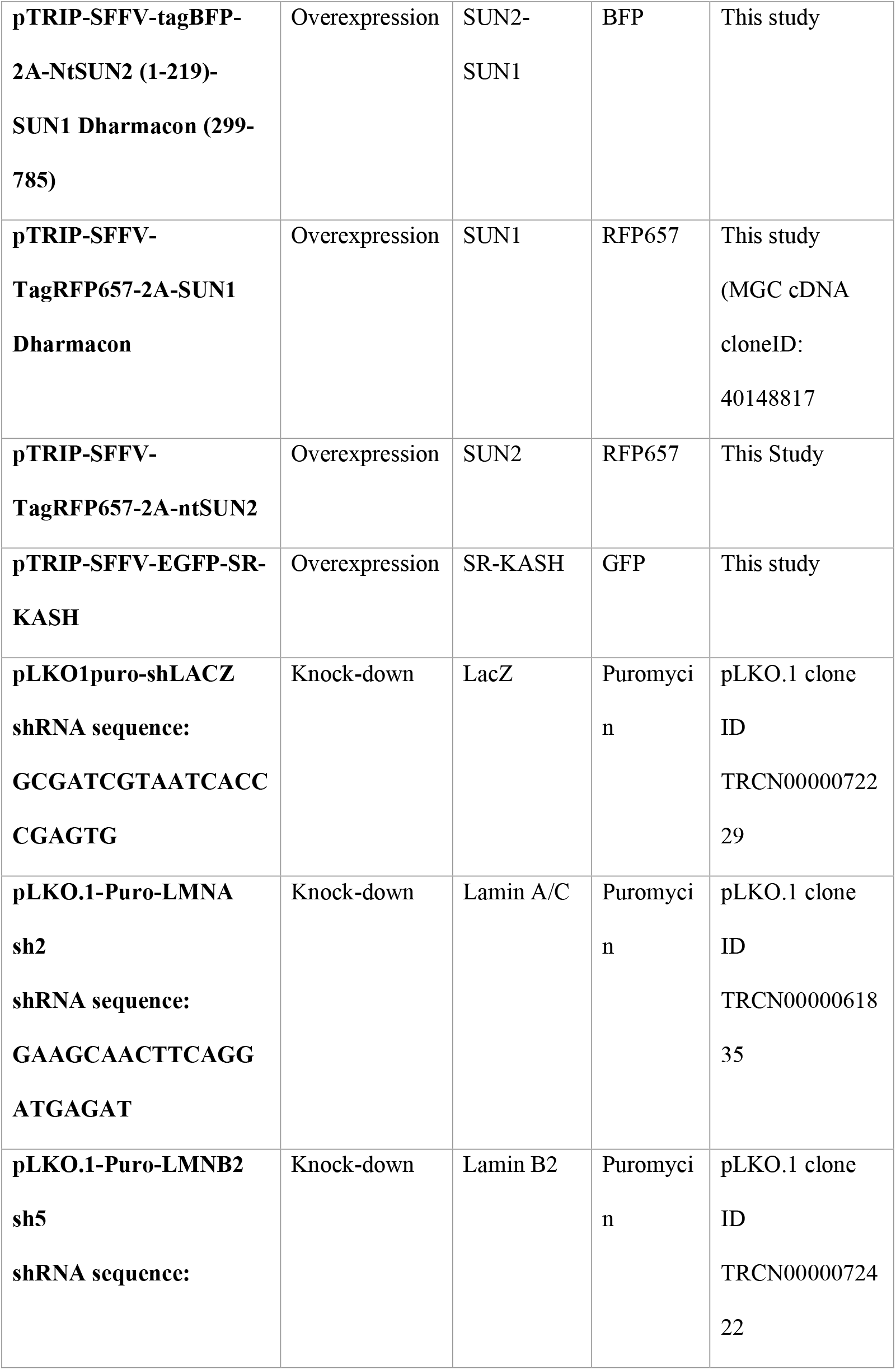
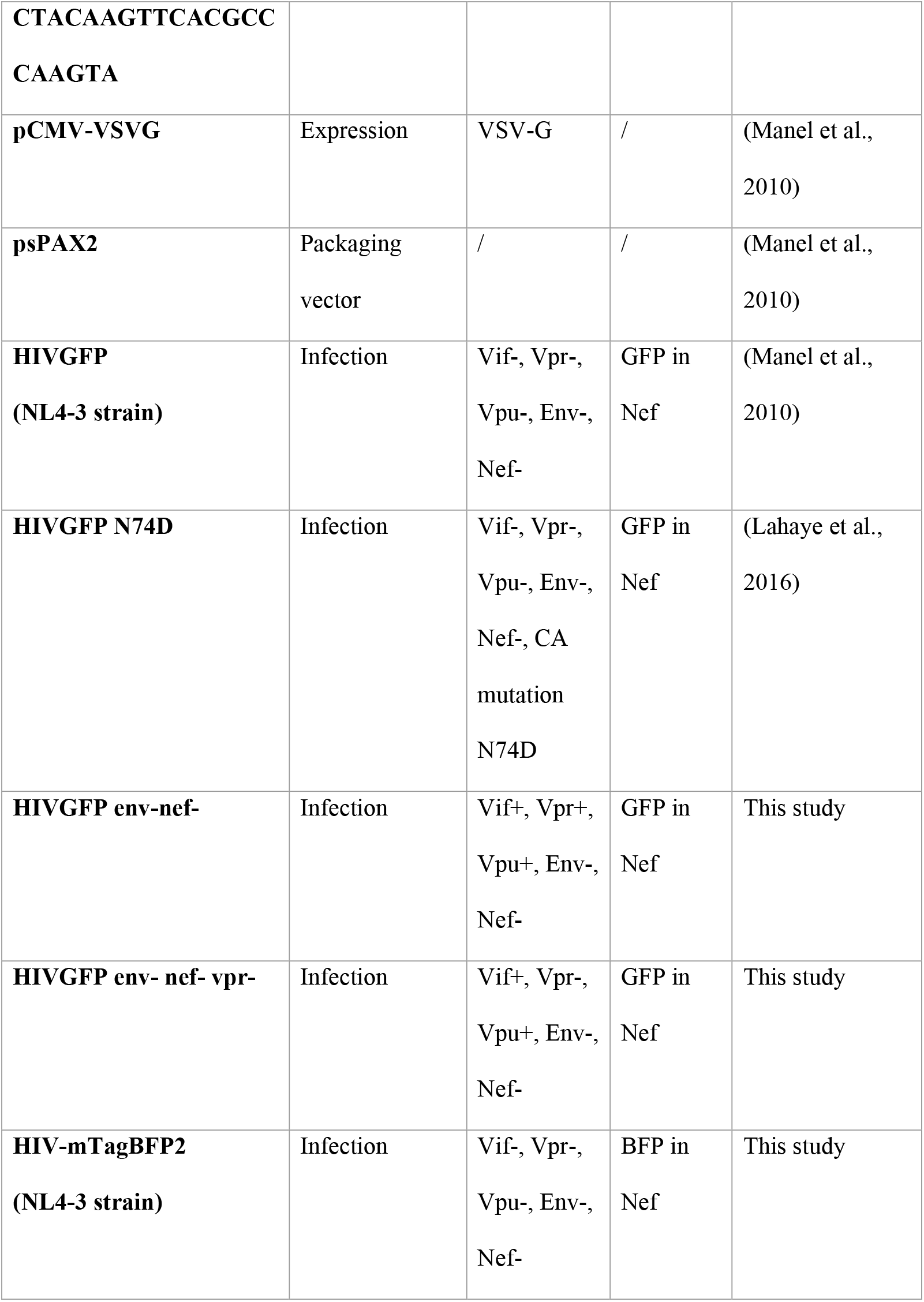
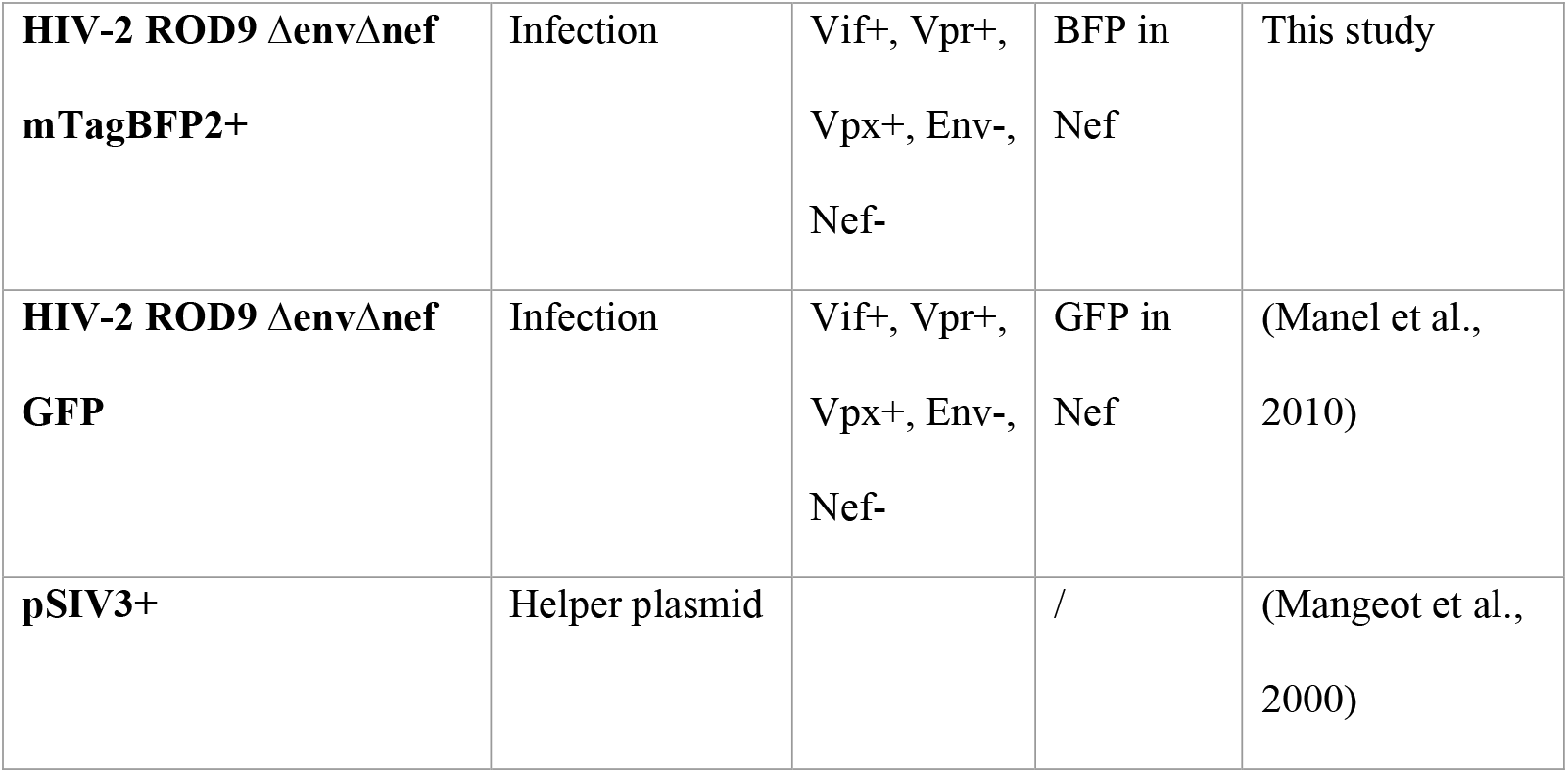
Plasmids used in this study.

**Table 2:** Drugs used in cell culture in this study.

| **Name** | **Description** | **Cat. Reference** | **Company** | **Final**<br>**Concentration** |
| --- | --- | --- | --- | --- |
| **Cyclosporin A** | Inhibits CypA-CA interaction | S2286 | Selleckchem | 2 μM |
| **AZT** | Reverse<br>transcriptase<br>inhibitor | A2169 | Sigma | 24 μM |
| **NVP** | Reverse<br>transcriptase<br>inhibitor | SML0097 | Sigma | 10 μM |
| **Etoposide** | Topoisomerase<br>II inhibitor | E1383 | Sigma | 5, 50 or 500 μM |
| **Q-VD-Oph** | Pan-caspase<br>inhibitor | S7311 | Selleckchem | 50 μM |
| **AZD6738** | ATR inhibitor | S7693 | Selleckchem | 1 μM |
| **DMSO** | Diluent | BDH1115 | VWR<br>Chemicals | adjusted |

**Table 3:**
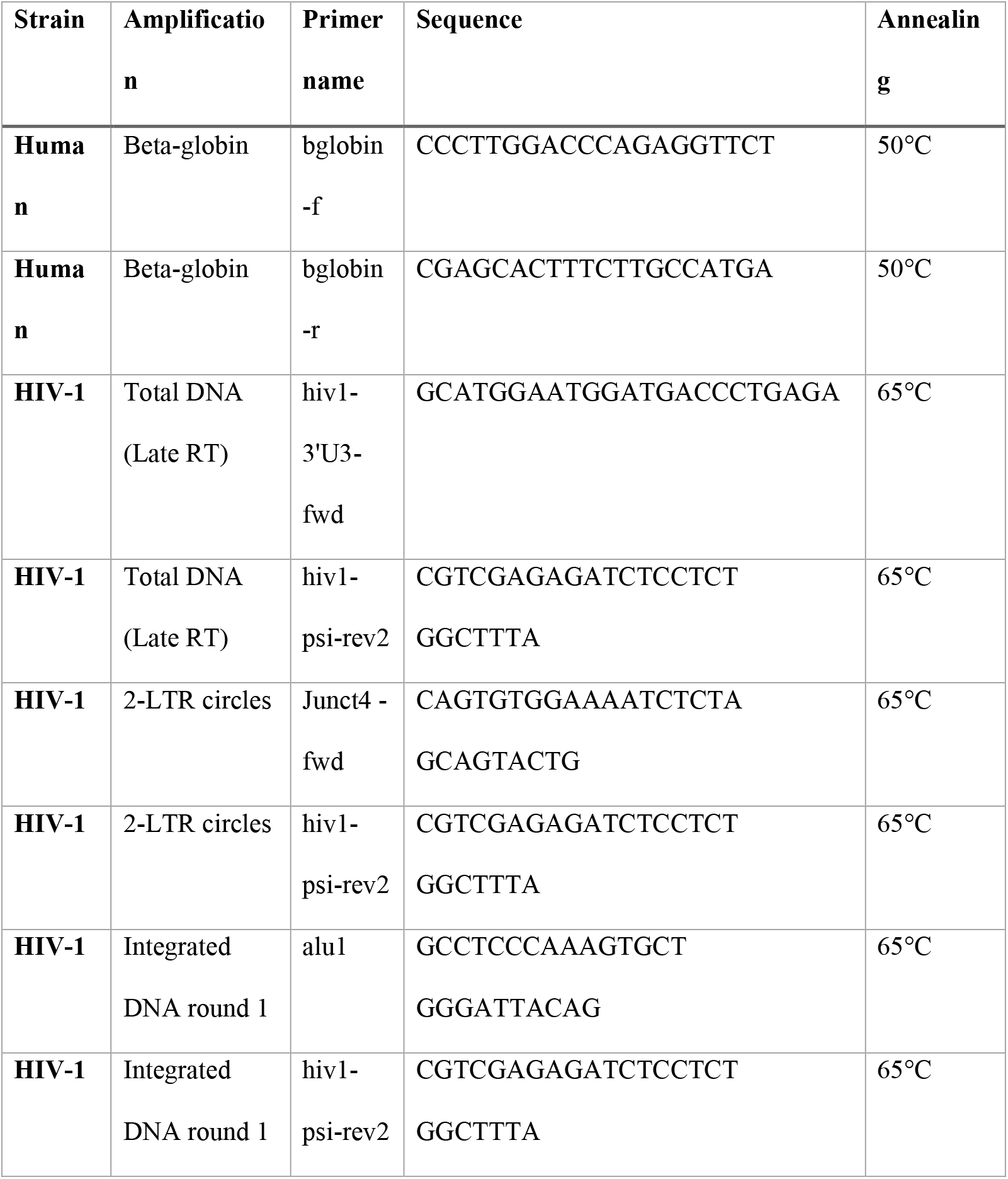
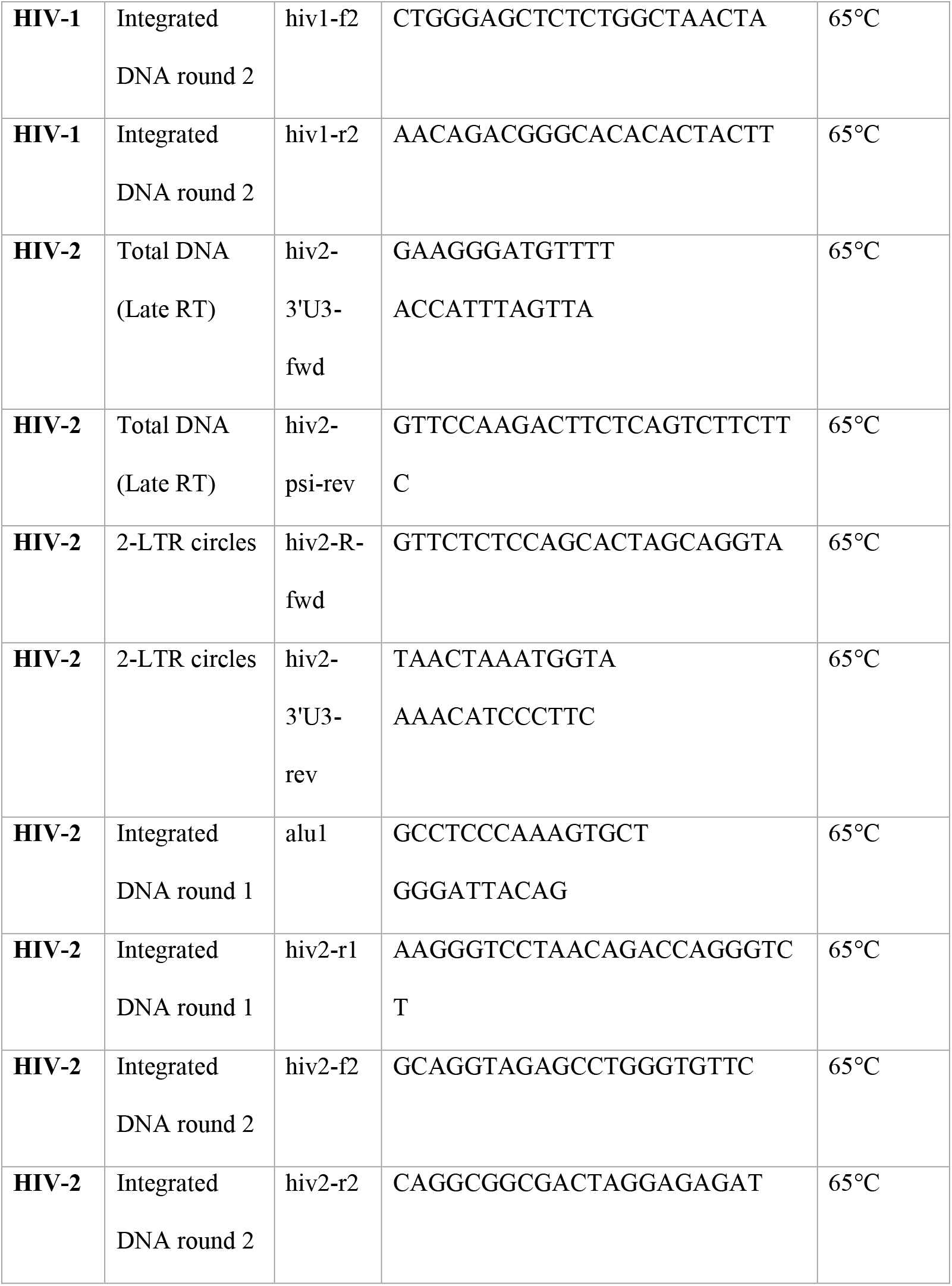
Primers used for HIV DNA species Real Time Quantitiative PCR.

**Table 4:**
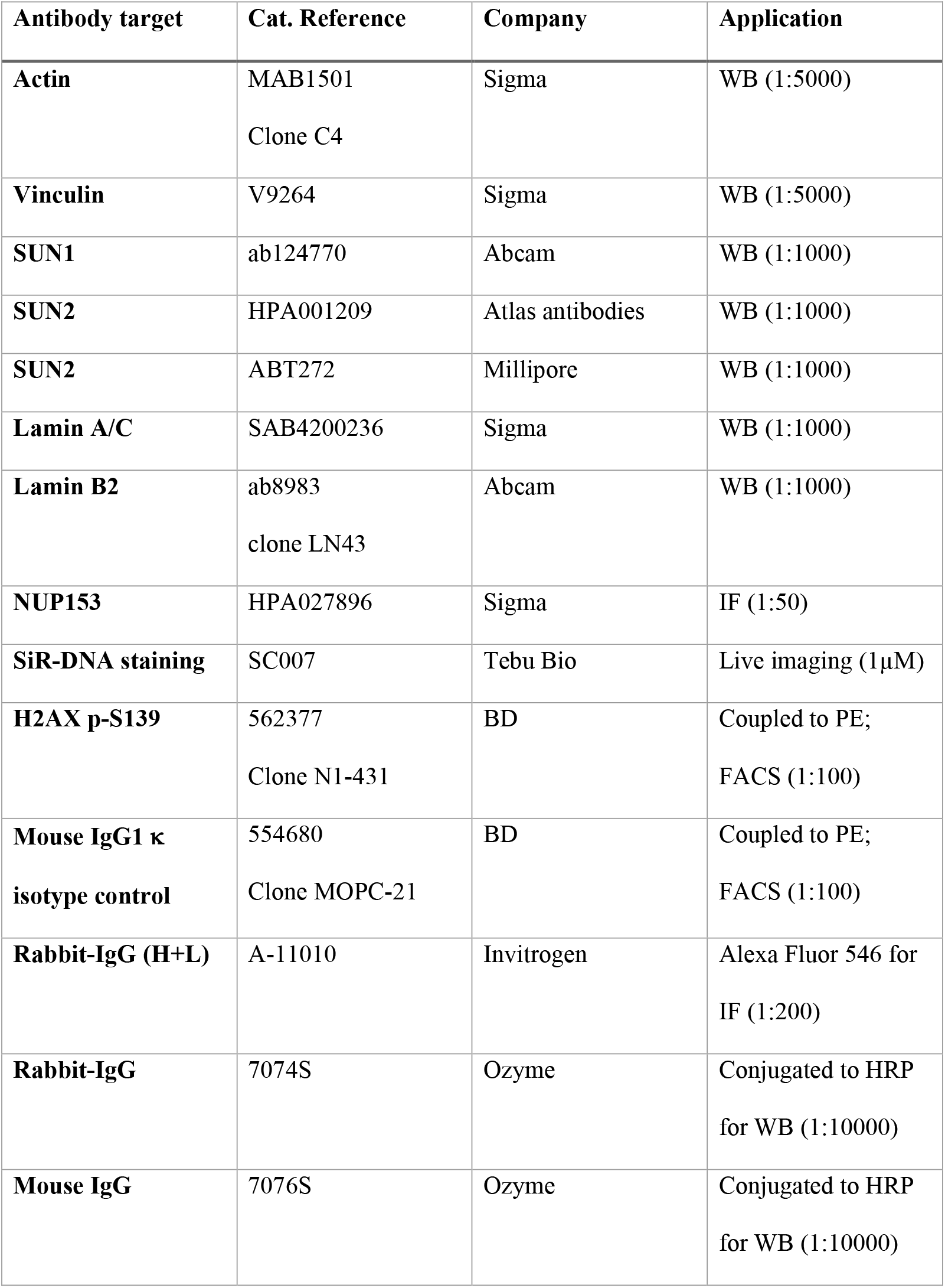
Antibodies used in this study for Western Blot, Confocal Imaging and Flow Cytometry.

